# DiSCs – Domains involving SETDB1 and Cohesin are critical regulators of genome topology and stem cell fate

**DOI:** 10.1101/2022.01.27.478011

**Authors:** Tushar Warrier, Chadi El Farran, Yingying Zeng, Benedict Shao Quan Ho, Don Loi Xu, Qiuye Bao, Zi Hao Zheng, Xuezhi Bi, Kelly Yu Sing Tan, Huck Hui Ng, Derrick Sek Tong Ong, Justin Jang Hann Chu, Amartya Sanyal, Melissa Jane Fullwood, James Collins, Hu Li, Jian Xu, Yuin-Han Loh

**Affiliations:** Cell Fate Engineering and Therapeutics Lab, Cell Biology and Therapies Division, A*STAR Institute of Molecular and Cell Biology, Singapore 138673, Singapore; Department of Biological Sciences, National University of Singapore, Singapore 117543, Singapore; Center for Individualized Medicine, Department of Molecular Pharmacology & Experimental Therapeutics, Mayo Clinic, Rochester, Minnesota 55905, USA; Proteomics Group, Bioprocessing Technology Institute, A*STAR, Singapore 138668, Singapore; Gene Regulation Laboratory, Genome Institute of Singapore, Singapore 138672, Singapore; School of Biological Sciences, Nanyang Technological University, 60 Nanyang Drive 637551, Singapore; Cancer Science Institute of Singapore, National University of Singapore, 14 Medical Drive, Singapore, 117599, Singapore; Howard Hughes Medical Institute, Boston, MA 02114, USA; Institute for Medical Engineering and Science Department of Biological Engineering, and Synthetic Biology Center, Massachusetts Institute of Technology, Cambridge, MA 02114, USA; Broad Institute of MIT and Harvard, Cambridge, MA 02139, USA; Wyss Institute for Biologically Inspired Engineering, Harvard University, Boston, MA, USA; Department of Plant Systems Physiology, Institute for Water and Wetland Research, Radboud University, Heyendaalseweg 135, 6525 AJ, Nijmegen, The Netherlands; NUS Graduate School for Integrative Sciences and Engineering, National University of Singapore, 28 Medical Drive, Singapore 117456, Singapore; Department of Physiology, Yong Loo Lin School of Medicine, National University of Singapore, Singapore 117593, Singapore; Department of Microbiology and Immunology, Yong Loo Lin School of Medicine, National University of Singapore, Singapore 117593, Singapore

**Keywords:** SETDB1, Cohesin, ChIP-Seq, H3K9me3-independent, pluripotency, topology, neuronal

## Abstract

SETDB1 is a key regulator of lineage-specific genes and endogenous retroviral elements (ERVs) through its deposition of repressive H3K9me3 mark. Apart from its H3K9me3 regulatory role, SETDB1 has seldom been studied in terms of its other potential regulatory roles. To investigate this, a genomic survey of SETDB1 binding in mouse embryonic stem cells across multiple libraries was conducted, leading to the unexpected discovery of regions bereft of common repressive histone marks (H3K9me3, H3K27me3). These regions were enriched with the CTCF motif that is often associated with the topological regulator Cohesin. Further profiling of these non-H3K9me3 regions led to the discovery of a cluster of non-repeat loci that were co-bound by SETDB1 and Cohesin. These regions, which we named DiSCs (Domains involving SETDB1 and Cohesin) were seen to be proximal to the gene promoters involved in embryonic stem cell pluripotency and lineage development. Importantly, it was found that SETDB1-Cohesin co-regulate target gene expression and genome topology at these DiSCs. Depletion of SETDB1 led to localized dysregulation of Cohesin binding thereby locally disrupting topological structures. Dysregulated gene expression trends revealed the importance of this cluster in ES cell maintenance as well as at gene ‘islands’ that drive differentiation to other lineages. The ‘unearthing’ of the DiSCs thus unravels a unique topological and transcriptional axis of control regulated chiefly by SETDB1.

## Introduction

Histone-lysine N-methyltransferase SETDB1 is an enzyme encoded by the *Setdb1* gene. It belongs to the Methyltransferase (EC 2.1.1) class of enzymes^1^. This protein chiefly mediates the tri-methylation of lysine residue 9 on histone H3 (H3K9me3). This trimethylation event constitutes an epigenetic silencing signal which recruits HP1 proteins to the methylated histones resulting in transcriptional repression^2,3^. Histone modifiers and transcription factors are widely seen to be critical regulators of cell fate and are also key determinants of cellular reprogramming^4,5^. SETDB1 also teams up with ATF7IP, which stabilises its methyltransferase activity^6,7^ and thereby allows it to repress endogenous retroviral elements (ERVs). SETDB1 is recruited by TRIM28, which is in turn recruited by KRAB zinc-finger proteins that can recognize specific sequences on ERVs, Long Terminal Repeats (LTRs) and other Transposable Elements (TEs)^8–10^. Among its other H3K9me3-associated functions, SETDB1 is also seen to pair up with OCT4 to promote the silencing of trophoblastic genes in mouse embryonic stem cells^11^. SETDB1 also associates with the PRC2 (Polycomb Repressive Complex 2) at developmental gene loci and contributes to their repression^12^. Multiple reports pertaining to the repressive role of SETDB1 encompass a variety of regulatory roles ranging from cell fate maintenance, genomic silencing and development^13^ and cellular reprogramming^14^. Recent studies have implicated other potential functions for SETDB1 ranging from proximal regulation of a large topological domain to methylation^15^ of non-histone substrate proteins in cancers^16–18^. Although the regulation of this proximal topological domain still pertains to the H3K9me3-specific role of SETDB1, its association to a topological region was fascinating. The non-histone associated roles of SETDB1 were revealed via identification of substrate-enzyme relationships in specific signalling cascades. An unbiased assessment of the overall binding profile of SETDB1 across the genome would thus serve as a preliminary clue towards deciphering its multiple roles in regulation.

Genome topology has two key regulatory components - CTCF (CCCTC-binding factor) and Cohesin^19^. CTCF is a transcription factor which contains 11 highly conserved zinc-finger domains^20^. Depending on its partners, CTCF can function as a transcriptional activator along with RNA Pol II^21^ or as a transcriptional repressor if bound to a histone deacytelase complex^22^. Additionally, CTCF demonstrates an insulating function between different compartments of the genome through the creation of Topologically Associating Domains (TADs)^23,24^. The Cohesin complex is deposited by NIPBL on to the genome, before it translocates along chromatin^25^. Cohesin and CTCF also play critical roles in dictating enhancer-promoter (E-P) contacts as a key component of topological regulation of the genome. This phenomenon has been seen to be particularly relevant to the maintenance of mouse embryonic stem cells (mESCs)^26^. Further studies have demonstrated additional roles for Cohesin complex in transcriptional regulation, DNA repair and chromatin condensation. Cohesin facilitates transcriptional insulation along with CTCF^27^. Studies have also shown that Cohesin is essential for stabilizing genomic loops^28,29^. The Cohesin complex binds in a dynamic manner across the genome wherein it passively slides across chromatin until it encounters CTCF. This sliding phenomenon engages the Cohesin complex in large loop formation^30,31^.

In this study, we discovered unique regions bound strongly by SETDB1, independent of repressive histone marks, which are co-bound with CTCF and Cohesin, with the enrichment for the latter being particularly strong. Further analysis of these sites, which were referred to as DiSCs (Domains involving SETDB1 and Cohesin) strikingly revealed their co-localization with activating histone marks such as H3K4me3 and H3K27Ac (promoter and enhancer associated marks). Using ChIP-Seq and RNA-Seq, we demonstrated that depletion of SETDB1 resulted in a drastic decrease of Cohesin binding at DiSCs, which was accompanied by the dysregulation of gene expression and genome topology locally. Hi-C data for mESCs revealed a strong enrichment of the DiSCs as topologically enriched regions. Ablation of SETDB1 and Cohesin further disrupted the topological structures at these DiSCs, led to gene compartment switches and contributed substantially to gene dysregulation. Interestingly, these dysregulated genes were key regulators of lineage-specific processes as well as maintenance of ES cell metabolic functions. This was in line with previously attributed functions of SETDB1 in ES cells – such as their survival, maintenance and for differentiation^32,33,34^. Detailed characterization of the DiSCs reveals a new mechanism of transcriptional and topological regulation by the SETDB1-Cohesin axis.

## Material and Methods

### Cell culture

E14 mouse embryonic stem cells (mESCs) were cultured using mouse ES medium prepared in DMEM high glucose (Hyclone) supplemented with 15% ES cell FBS (Gibco), 2mM L-glutamine (Gibco), 1X Pen-Strep (Gibco), 100 μM MEM non-essential amino acids (Gibco), 100μM β-mercaptoethanol (Gibco), and 1000 U/mL leukemia inhibitory factor (LIF; ESGRO, Millipore). The mouse ES line was cultured on plates pre-coated with 0.1% gelatine (porcine). Mouse embryonic fibroblasts (mEFs) and 293T cells were cultured directly on non-coated, treated culture plates. The media for mEF cells was prepared using DMEM high glucose (Hyclone), along with 10% Heat Inactivated (HI) FBS, 2mM L-Glutamine (Gibco), 1X Pen-Strep (Gibco) and 100 μM MEM non-essential amino acids. All cultures were maintained at 37°C with 5% CO2.

### Plasmid construction

We used the pGL3-promoter vector with a SV40 promoter upstream of the luciferase gene for cloning DiSC fragments for the Luciferase assays. For the shRNA constructs, we utilized the pSUPER.puro vector for cloning the shRNA oligos against *Setdb1, Smc1a, Atf7ip* and *Sumo2*. For the CRISPR-Cas9 experiments, we used the lentiCRISPR v2 (Addgene plasmid #52961) to clone the sgRNA sequences. For the constructs used in over-expression and coimmunoprecipitation experiments, we cloned our ORF sequences (*Setdb1, Smc1a*) into pCAG-HA-puro, pCAG-FLAG-puro and pSIN.

### shRNA design

Dharmacon’s siDESIGN center (http://dharmacon.gelifesciences.com/design-center/) was used for the design of the shRNA sequences. shRNA sequences were ordered as DNA oligos from IDT and cloned into the pSUPER.puro plasmid (**Supplementary Table S1**).

### Transfection into mESCs

E14 cells were harvested using trypsinization and were re-seeded into 6-well plates which were pre-coated with 0.1% gelatine. The seeding density was 300,000 cells per well of the plates. These cells were cultured at 37°C for 20-24 hours and the cells were replenished with fresh medium at least 2 hours prior to transfection. The cells were then transfected with 3 μg of the plasmid constructs using 4.5 μL of Lipofectamine 2000 (Thermofisher), as per the manufacturer’s protocol. Puromycin (1 μg/mL) containing mESC medium was added to the cells 24 hours after transfection to select for the transfected plasmids. This process of selection was repeated once more 48 hours after transfection. The cells were finally harvested 72 hours after transfection.

### Chromatin Immunoprecipitation

For preparation of chromatin the protocol previously described^35^ was utilized. Cells were trypsinized, harvested and the cell number was estimated. Cross-linking was performed using 1% formaldehyde for 10 min at room temperature followed by quenching with 0.125 M Glycine. Cross-linked pellets were washed twice with ice-cold 1X PBS (with 0.1% Triton X-100) and then subjected to lysis with a lysis buffer (10 mM Tris-Cl (pH 8), 100 mM NaCl, 10 mM EDTA, 0.25% Triton X-100 and protease inhibitor cocktail (Roche)). Pellets were resuspended after centrifugation, in 1% SDS lysis buffer (50 mM HEPES-KOH (pH 7.5), 150 mM NaCl, 1% SDS, 2 mM EDTA, 1% Triton X-100, 0.1% NaDOC and protease inhibitor cocktail). Following complete resuspension, the samples were nutated at 4°C for 15 mins and the pellets were re-obtained after high-speed centrifugation. This was followed by two washes of all the lysed samples with 0.1% SDS lysis buffer (50 mM HEPES-KOH (pH 7.5), 150 mM NaCl, 0.1% SDS, 2 mM EDTA, 1% Triton X-100, 0.1% NaDOC and protease inhibitor cocktail). For every 10 million cells, 14 cycles of sonication were carried out using the Bioruptor (Diagenode) with 30 sec pulses and 60 sec halts in each cycle. Cell debris was separated from the sheared chromatin by centrifugation at 15,000 rpm at 4°C for 30 mins. Preclearing of the chromatin was carried out with 100 uL of Protein G Dynabeads (Life Technologies) for 2 hours at 4°C. Simultaneously, 100 uL of Protein G Dynabeads were also bound to 5 ug (per 10 million cells) of the antibodies. After separation of a small amount as input - the remaining pre-cleared chromatin was used to bind to the antibody-bound beads at 4°C, overnight. Elution involved three washes with 0.1% SDS lysis buffer, one with 0.1% SDS lysis buffer/0.35 M NaCl, one with 10 mM Tris-Cl (pH 8.0), 1 mM EDTA, 0.5% NP40, 0.25 LiCl, 0.5% NaDOC, and one with TE buffer (pH 8.0). The immunoprecipitated chromatin was eluted out from the beads by heating the beads resuspended in 50 mM Tris-HCl (pH 7.5), 10 mM EDTA, 1% SDS, for 1 hour at 68°C while shaking at 1400 rpm. Cross-links were reversed by incubating the eluted samples and inputs at 42°C for 2 hours and 67°C for 6 hours in the presence of Pronase (Sigma) and TE buffer, after which the DNA was purified using the QIAGEN PCR Purification kit (for inputs) and the QIAGEN MinELute PCR Purification kit (for samples) as per the manufacturer’s instructions. Quantitative PCR was performed for the purified ChIP-DNA samples by using the CXF384 Real-time System (Bio-Rad), using a Kapa SYBR Fast qPCR kit (Kapa Biosystems). The data represented was normalized to the Inputs as well as the Negative control primers, thus representing an overall fold enrichment.

### Sequential Chromatin Immunoprecipitation (Sequential ChIP)

Protein A and Protein G Dynabeads were combined in an equimolar ratio (50 μL per sample) and washed thrice with 1X PBS (with 0.1% Triton X-100). The beads were then resuspended in 600 μL of Pre-Adsorption buffer (equal amounts of ChIP lysis buffer and ChIP dilution buffer (1% Triton X-100, 2mM EDTA, 20 mM Tris-HCl, 150 mM NaCl, 1X Protease inhibitor cocktail)). BSA was added to the resuspended beads at a final concentration of 200 μg/mL. This process of pre-adsorption and blocking of the beads to allow specificity in the pull-down was carried out overnight at 4°C. After pre-adsorption, the beads were washed with ChIP dilution buffer and the first antibody (either SMC1A or CTCF) to 100 μL of the bead mixture. The antibody was bound to the beads at room temperature over a period of 3 hours. Sonicated E14 DNA (see Chromatin Immunoprecipitation protocol) was pre-cleared with 100 μL of the beads for 3 hours at 4°C. After pre-clearing, 50 μL of the chromatin was separated out as input and the rest was added to the antibody-bound beads and then incubated at 4°C, overnight. For the first elution, the beads were washed thrice with 0.1% SDS lysis buffer, once with 0.1% SDS lysis buffer/0.35 M NaCl, once with 10 mM Tris-Cl (pH 8), 1 mM EDTA, 0.5% NP40, 0.25 LiCl, 0.5% NaDOC, and once with TE buffer (pH 8.0). The beads were separated by a short spin at 800g for 1 minute. The beads were resuspended in 75 μL of elution buffer (50 mM Tris-HCl (pH 7.5), 10 mM EDTA, 1% SDS) and kept at 37°C for 30 minutes. The DNA was eluted from the beads by a brief spin at 1000g for 2 minutes. 15 μL of the eluted sample was separated out for validation of the first ChIP. The remaining 60 μL was diluted with ChIP dilution buffer to 1200 μL.

The second antibody (SETDB1 or SMC1A) was added to freshly pre-adsorbed 100 μL Protein A and G Dynabeads and allowed to bind at room temperature for 3 hours. In order to maintain a 1:19 Input:IP ratio, 63 μL of the first eluted sample was separated out. The rest of the sample was added to the antibody-bound beads and incubated overnight at 4°C. The elution steps for the second ChIP were the same as the first one and then eluted at 68°C for 60 minutes while shaking at 1400 rpm. Following, this the samples were de-crosslinked, purified and subsequently used for qPCR.

### ChIP-Seq library preparation

ChIP samples and the corresponding inputs were prepared as mentioned before (**See Chromatin Immunoprecipitation in Methods**). Following qPCR validation of the samples, libraries were prepared by using Illumina reagents with TruSeq adapters as per the manufacturer’s instructions. The quality and concentration of each sample was assessed by using Agilent High Sensitivity chips on the Agilent 2100 Bioanalyzer. High throughput sequencing for the samples was performed on the HiSeq 4000 (Illumina).

### ChIP-Seq Analysis

Sequenced ChIP-Seq libraries were first run through the FastQC software (https://www.bioinformatics.babraham.ac.uk/projects/fastqc/) to determine the quality of the libraries. The libraries were then mapped to the mm9 genome assembly by using the STAR aligner while setting the –alignIntronMax option to 1 and the –alignEndsType option to EndToEnd to ensure compatibility of the ChIP-Seq files to STAR. Published library datasets were mapped in a similar manner. To include repeat regions and multi-mapped regions of the genome, we set the –outFilterMultimapNmax option to 500. The makeTagDirectory script of HOMER was executed while using the -keepOne option to retain the repeat elements that were mapped to the genome. Peaks were called for the libraries by using the findPeaks script of HOMER with the -style being set to factor for proteins such as CTCF. For the histone marks, the -style was set to histone. The same was done for the detection of the broad peak binding profile of SETDB1 and SMC1A in the study. All the discovered peaks were annotated by using the annotatePeaks.pl script. For visualization of the ChIP-Seq libraries we generated UCSC bedgraph files by using the makeUCSCfile script on HOMER. The following libraries were downloaded for analysis from the Gene Expression Omnibus (GEO)^36^: E14-SETDB1 (GSM440256)^8^, V6.5-SETDB1 (GSM459273, GSM459274, GSM459275)^32^, H3K9me3 (GSM440257)^11^, H3K27me3 (GSM1327220)^37^, H3K27Ac (GSM1000126)^38^, H3K9Ac (GSM1000123)^39^, H3K4me3 (GSM1000124)^39^, CTCF (GSM699165)^21^, NIPBL (GSM560349)^40^, SMC3 (GSM560344)^41^, OCT4 (GSM288346)^42^, NANOG (GSM288345)^42^, SUMO2 (GSM1819197)^43^.

### RNA-Sequencing

Total RNA was extracted for each of the transfected cells using Trizol reagent (Ambion). DNA contamination for the samples was minimized by using the QIAGEN RNeasy Kit. The RNA samples were processed using a TruSeq Stranded mRNA Library Prep Kit (RS-122-2101, Illumina). This kit was used for mRNA selection, fragmentation, cDNA synthesis and library preparation. The library quality was analyzed on an Agilent Bioanalyzer using the Agilent DNA 1000 kit. High throughput sequencing was then performed on a HiSeq4000 instrument.

### RNA-Seq Analysis

For the RNA-Seq analysis, the quality control of the libraries was performed by using the RSeQC package^44^ and SeqMonk (https://www.bioinformatics.babytheraham.ac.uk/projects/seqmonk/). The libraries were mapped to the mm9 genome assembly by using the STAR aligner^45^. All reads that were mapped to more than one genomic locus or had multiple mismatches were filtered out. Gene expression levels and differential gene expression were assessed by using the cuffdiff tool^46^. For visualizing the RNA-Seq libraries on the UCSC browser, the bedgraph files were generated by using the makeUCSCfile script with the addition of the option -style rnaseq. For the gene lists, all genes below FPKM values of 0.5 were filtered out.

### Chromosome conformation capture (3C)

As per a recommended^47^, 40 million cells were harvested and washed twice with ice-cold PBS. Cells were then cross-linked by using 1% Formaldehyde (Sigma) for 10 minutes at room temperature. The reaction was quenched with 0.266 M of Glycine followed by washing twice with ice-cold PBS. The pellets were lysed on ice for 15 min after resuspending in 1 mL of 3C lysis buffer (10 mM Tris–HCl (pH 7.5), 10 mM NaCl, 5 mM EDTA, 0.5% NP-40, 1X Protease inhibitor). The lysates were further dounced by using a tight and loose douncer, to ensure effective lysis. Nuclei were pelleted by centrifugation and were resuspended in 500 μL of NEB Buffer 3.1 (1X) and washed twice in this buffer and then split into 20 tubes with 25 μL each and spun down. After this, 362 μL of 1X Buffer 3.1, 33 μL of 1% SDS were added to each tube and then incubated at 65°C, 10 minutes. This was followed by addition of 44 μL of 20% Triton X-100 (Sigma-Aldrich) and 200 U of BglII for enzymatic digestion overnight. Some of the lysate was separated out before and after digestion for using as a control to estimate digestion efficiency by qPCR. The digested DNA was pooled together and treated with the ligation mix (40 mL nuclease-free water, 5 mL 10X Ligation Buffer, 2.687 mL 20% Triton X-100) for 1 hour at 37°C. After cooling the mix on ice for 10 minutes, 67 kilo-units (KU) of NEB T4 DNA Ligase was added and incubated overnight at 16°C. 15 μL of 20 mg/mL Proteinase K (Promega) was then added to the mixture and incubated at 65°C overnight. The samples were cooled to room temperature and 30 μL of 10 mg/mL RNase A (QIAGEN) was added and the samples were incubated at 37°C for 1 hour. The 3C libraries were purified by phenol-chloroform extraction and PCR purification kits (QIAGEN). The 3C interactions were then detected by qPCR with the use of a Kapa SYBR Fast qPCR kit.

### Co-Immunoprecipitation

pCAG-FLAG-*Smc1a* and pSIN-V5-*Setdb1* were co-transfected into 293T cells by using Lipofectamine 2000. These cells were cultured for a period of 72 hours after transfection and then harvested and lysed by using the lysis buffer (50 mM Tris-HCl, pH 8.0, 150 mM NaCl, 5 mM EDTA, 0.5% NP-40, 1X protease inhibitor cocktail). Immunoprecipitation (IP) was carried out for FLAG-SMC1A by using 100 μL of Protein A Dynabeads, overnight at 4°C, and then for V5-SETDB1. For elution the beads were washed thrice with the lysis buffer and then resuspended in 2X Laemmli sample buffer (BioRad) and boiled for 10 minutes, prior to loading on to an 8% SDS-PAGE gel.

### Western Blot

Cells were lysed by using lysis buffer (containing protease inhibitor cocktail and PMSF) and the lysate was separated at 12,000 g, 20 minutes. As per the requirement, the concentrations of the protein samples were estimated by Bradford assay. The proteins in the lysate are denatured by boiling for 10 minutes with 2X Laemmli buffer. The protein samples were loaded onto an SDS–PAGE gel and transferred onto a polyvinylidene difluoride membrane (BioRad). The membrane was blocked with 3% BSA at room temperature overnight, followed by incubation with primary antibodies for 1.5 hours at room temperature. The secondary horseradish peroxidase (HRP)-conjugated anti-mouse IgG, HRP-conjugated anti-rabbit IgG or HRP-conjugated anti-goat IgG (1:10,000) antibodies were then added to the membrane at room temperature for 1 hour. For signal detection, we used the SuperSignal West Dura Extended Duration Substrate (Thermo Scientific) and captured the blots on CLXposure films (Thermo Scientific). The primary antibodies used were anti-V5 (1:1000, ThermoFisher Scientific #R960-25), anti-ACTIN (1:1000, Santa Cruz #sc-10731), anti-FLAG (1:500, Sigma-Aldrich #F1804), anti-ESET (SETDB1) (1:2000, Santa Cruz #sc-66884), anti-SMC1A (1:2000, Bethyl #A300-055A) and anti-H3K9me3 (1:1000, Abcam #ab8898).

### Mass Spectrometry

E14 cells were cultured in three 15-cm Corning treated, cell culture dishes. After culturing for 72 hours, the cells were harvested and lysed by using lysis buffer. SMC1A antibody (Bethyl, A300-055A) and SETDB1 antibody (home-made) were incubated with 100 μL of Protein A Dynabeads for 3 hours at room temperature. The lysate was then added to this mix and incubated overnight. The protein was eluted out of the antibody-bead complex by washing 3 times with lysis buffer, followed by boiling for 30 minutes with Elution buffer 1 (20 mM citrate acid, pH 6 + 0.1% Tween-20). This was followed by another round of elution with Elution buffer 2 (100 mM TEAB, pH 8.5 in 10% SDS). Peptides were extracted and evaporated to dryness in SpeedVac (Savant Instruments, Holbrook, NY, USA), and then dissolved in LC-MS buffer: 2% (v/v) acetonitrile-1% (v/v) formic acid. Mass spectrometry analysis was performed on LTQ-Orbitrap Elite Mass Spectrometer (Thermo Fisher Scientific, San Jose, CA), which was equipped with a nanoACQUITY UPLC system (Waters Milford, MA), Thermo Xcalibur 3.063 and LTQ Tune Plus 2.7.0.1112 SP2 instrument control. Mobile phases composed of A: 0.1% (v/v) formic acid in water, and B: 0.1% (v/v) formic acid in acetonitrile. Peptides (5 μL) were desalted on Symmetry C18 trapping column, 5 μm, 180 μm x 20mm (Waters) for 5 minutes with a 1% mobile phase B at 8 μL/min. The desalted peptides were separated on nanoACQUITY UPLC BEH130 C18 column, 1.7 μm, 75 μm x 200 mm, over a 90-minute linear gradient from 5% to 40% mobile phase B. Ionization was achieved by nano-spray in the positive ion mode at 1.8 kV. Spectra were obtained by data-dependant scanning tandem MS, in which one full MS scan at 120,000 resolution from 350 to 1,600 m/z was followed by HCD Orbitrap tandem MS scans of the 15 most intense peptide ions, fragmented with normalized collision energy of 35.0%, at a resolution of 15,000.

### Hi-C Seq

Cells treated with the shControl, sh*Setdb1* and sh*Smc1a* constructs were harvested at the 72-hour time point post transfection. Assessment for the depletion levels of the *Setdb1* and *Smc1a* transcripts was performed and seen to be >75% depleted. The cells were fixed using 2% formaldehyde and subsequently quenched with 0.125M glycine. The amount of input DNA per million cells of each type was determined using the protocol recommended by the Arima-HiC+ kit. Using the recommended kit, the cells equivalent to 3-5 ug of DNA were used as starting material and were lysed. The DNA released was RE digested, ends were filled-in using biotin and then ligated. The proximally ligated DNA was then sheared using a Covaris sonicator in the range of 300-700 bp and was enriched for biotin-bound DNA prior to ligation to indexed adapters. Finally, the library was amplified and sequenced using an Illumina HiSeq4000 instrument with one sample sequenced across 3 lanes for sufficient depth.

### Quantitative PCR

For cDNA samples to be quantitated by qPCR, RNA was converted to cDNA by using the 5X iScript Reverse Transcriptase mix (BioRad). The cDNA samples, after dilution, were run on the CXF384 Real-time System (Bio-Rad), using a Kapa SYBR Fast qPCR kit (Kapa Biosystems). *Actin* or *Gapdh* were used as control primers to normalize gene expression. For ChIP DNA samples, negative control primers were designed based on lack of binding of factors (gene desert regions).

### Luciferase assay

cDiSC and ncDiSC fragments were cloned into the pGL3-promoter vector flanked by the KpnI and XhoI restriction sites. E14 cells were co-transfected with pGL3-promoter vector clones and pRL Renilla luciferase control vector using Lipofectamine 2000. Cells were harvested 72 hours post-transfection using the Passive lysis buffer (provided with the Dual Luciferase Reporter Assay system (Promega)). Luciferase assay was performed as per the the GloMAX Explorer system protocol.

### sgRNA design

Sequences of the SETDB1 and Cohesin binding sites at DiSCs were imported into the CRISPR sgRNA designer^48^ and the outputs of the same were checked for their off-target scores. The sgRNA sequences with the lowest off-target scores were chosen and then cloned into the lentiCRISPR v2 vector and the sgRNA constructs were transfected into the E14 cells by using Lipofectamine 2000.

### SURVEYOR assay

E14 cells were transfected with the cloned sgRNA constructs using Lipofectamine 2000 and the transfected cells were then selected using Puromycin (1 μg/mL) from 24 hours post-transfection. After 2 rounds of selection, the cells were harvested 72 hours after transfection. Genomic DNA was extracted from the harvested cells using QIAGEN DNeasy Blood & Tissue Kit following manufacturer’s protocol. The CRISPR target sites for each of the sgRNA was PCR amplified and purified using QIAquick PCR Purification Kit following manufacturer’s protocol. Purified PCR products (400 ng) were mixed with 2 μL 10X Taq polymerase PCR buffer (QIAGEN) and nuclease free water to a final volume of 20 μL. The PCR products were annealed: 95°C for 10min, 95°C to 25°C (ramping at – 0.25°C/s). After re-annealing, products were treated with SURVEYOR nuclease and SURVEYOR enhancer (IDT), following the manufacturer’s recommended protocol. These products were analyzed on a 2% agarose gel and was imaged with Gel Doc imaging system (BioRad). The observation of multiple bands implied successful targeting of the sequences by the sgRNA.

### Construction of FLAG-*Setdb1* cell line (C1)

To generate the 3XFLAG-*Setdb1* knock-in cell line, we designed sgRNA targeting the upstream 5’ region of the *Setdb1* gene using the CRISPR sgRNA designer. This sgRNA was cloned into lentiCRISPR v2. A donor plasmid pCAG-puro was used for cloning the homologous DNA sequence for *Setdb1* with 3XFLAG. The CRISPR and donor plasmids were co-transfected into E14 cells and two rounds of puromycin selection were carried out. The colonies arising out of a single cell were used for immunostaining with FLAG and further confirmed by Sanger sequencing. The confirmed colonies were then expanded to attain a stable cell line.

### Immunofluorescence

Cells were fixed for 30 minutes at room temperature using 4% Paraformaldehyde solution. This was followed by permeabilization using 0.25% Triton X-100 for 15 minutes. Blocking was performed by using 1% BSA/0.2% Tween 20 solution for 30 minutes. The cells were then treated with the anti-FLAG (Sigma, F1804) antibody followed by the Alexa 488 conjugated anti-mouse secondary antibody (Thermofisher) and counter-stained with Hoechst 33342 (Thermofisher). A Zeiss LSM700 confocal microscope was used for capturing the images.

### Knock down Hi-C analysis

The Hi-C reads were aligned to the mm9 genome using BWA with runHiC pipeline using the default settings and set the enzyme as Arima (https://pypi.org/project/runHiC/). Valid pairs were retained and output as .cool file which used for downstream analysis. Cooler file were input to the HiCPeaks (https://github.com/XiaoTaoWang/HiCPeaks) with BH-FDR algorithm for loop detection using 5kb and 10kb resolution^49^. Differential compartments were called using dcHiC using 100kb resolution (https://github.com/ay-lab/dcHiC)^50^. Pile-up analysis of the loops were performed by coolpup.pl using the .cool file (https://github.com/open2c/coolpuppy)^51^. The genes enriched in differential compartments were defined by the overlap between the gene regions and differential compartments regions using bedtools.

### Cohesin Hi-ChIP analysis

Cohesin HiChIP paired-end reads in mESC (GSE80820)^52^ were aligned to the mm9 genome using HiC-Pro software^53^. Reads were assigned to the MboI restriction enzyme fragments and duplicates were removed from analysis. Valid interaction pairs were generated and converted to the .hic file using Juicer tools^54^. HiCCUPs^49^ from the Juicer package was used to call the significant loops and the Knight-Ruiz (KR) normalization method^55^ was used for the correction of the contact matrix. And the parameters used for calling loops were set as follows: -m 500 -r 5000,10000 -f 0.1,0.1 -p 4,2 -i 7,5 -d 20000,20000. DiSC genomic regions were overlapped with loop anchors by using bedtools, with a threshold setting of minimal 1bp overlap. Finally, aggregate peak analysis (APA)^49^ was used to evaluate the enrichment of putative SMC1A loops on the DiSC.

### Differential ChIP-Seq Analysis

By using the getDifferentialPeaks function for Homer, we were able to identify differential peaks between shControl Smc1a and sh*Setdb1* Smc1a ChIP-Seq libraries. The .bed file supplied for this contained either the cDiSC or ncDiSC co-ordinates to obtain the differential peaks specifically from these subsets. The fold change option -F was set to 2 to identify the sites with the most significant effects.

### Reprogramming

Immortalized mEF (imEF) cells were cultured on the Corning cell culture plates. These cultured imEF cells (2000) were seeded onto Corning 12-well plates and infected with OSKM virus simultaneously. We used a feeder free reprogramming medium (DMEM/F12 (Gibco) supplemented with 15% Knockout Serum Replacement (Gibco), 2 mM L-glutamine (Gibco), 1× Pen-Strep (Gibco) 100 μM MEM non-essential amino acids (Gibco), 100 μM β-mercaptoethanol (Gibco), and 1000 U/mL LIF (ESGRO, Millipore), 50 μg/mL ascorbic acid (Sigma), 10 ng/mL basic fibroblast growth factor (bFGF, Gibco), 3 μM CHIR99021 (Stemgent) and 0.5× N-2 Supplement (Gibco)) which was changed daily for the cells. After 16 days of reprogramming, the iPSC-like cells were harvested and cross-linked and subsequently utilized for ChIP using the SETDB1 and SMC1A antibodies.

### Enrichment heatmaps

The deeptools software package was utilised for plotting heatmaps based on the ChIP-Seq data. The bamCoverage function was used to create bigwigs out of the mapped files and normalization was performed using the RPGC option with the -effectiveGenomeSize being set for the mm9 assembly. A matrix was generated using the computeMatrix function with the scale-regions and --skipZeros options being utilized. Finally, heatmaps were generated by using the plotHeatmap function.

### Clustering Analysis of ChIP-Seq libraries

Clustering analysis of ChIP-Seq libraries was performed by first generating Jaccard indices of the bound loci. This was calculated by using BEDTools^56^. The Jaccard indices were then clustered by using R for the generation of heatmaps.

### Gene Ontology Analysis

For performing gene ontology analysis, any duplicates in the gene list of interest were removed and the filtered list was input into Metascape (http://metascape.org)^57^, while selecting *Mus musculus* as the input species. The list of gene ontology terms was then ordered by the [-log(p-value)] for representation.

### Interaction Networks

The gene names were input into the STRING database^58^ and the protein-protein interaction networks were formulated by setting the stringency to low (score = 0.150) and filtering out any text-mined interactions. The interaction networks were saved as .tsv files. For determining relevant sub-networks within the large networks, we input the list of interacting proteins into the MCODE^59^ tool within Cytoscape^60^.

## Results

### SETDB1 and Cohesin co-bind to DiSCs devoid of repressive histone marks

To assess the genome-wide binding profile of SETDB1 across ES (embryonic stem) cells, we prepared SETDB1 ChIP-Seq libraries in the E14 mouse embryonic stem cell (mESC) line. SETDB1 is known to be associated with the repressive histone marks H3K9me3 and H3K27me3, which are required for endogenous retroviral (ERV) element silencing in ES cells^61,62^. Our ChIP-Seq analysis revealed that a significant proportion (~40%) of SETDB1-bound sites (cluster 1) were devoid of the repressive H3K9me3 epigenetic marks (**Figure 1A**, **Supplementary Figure S1A**). The cluster 1 SETDB1 and SMC1A co-bound sites were also completely free from H3K9me3 binding (**Supplementary Figure S1B, Supplementary Figure S1C**).

**Figure 1.**
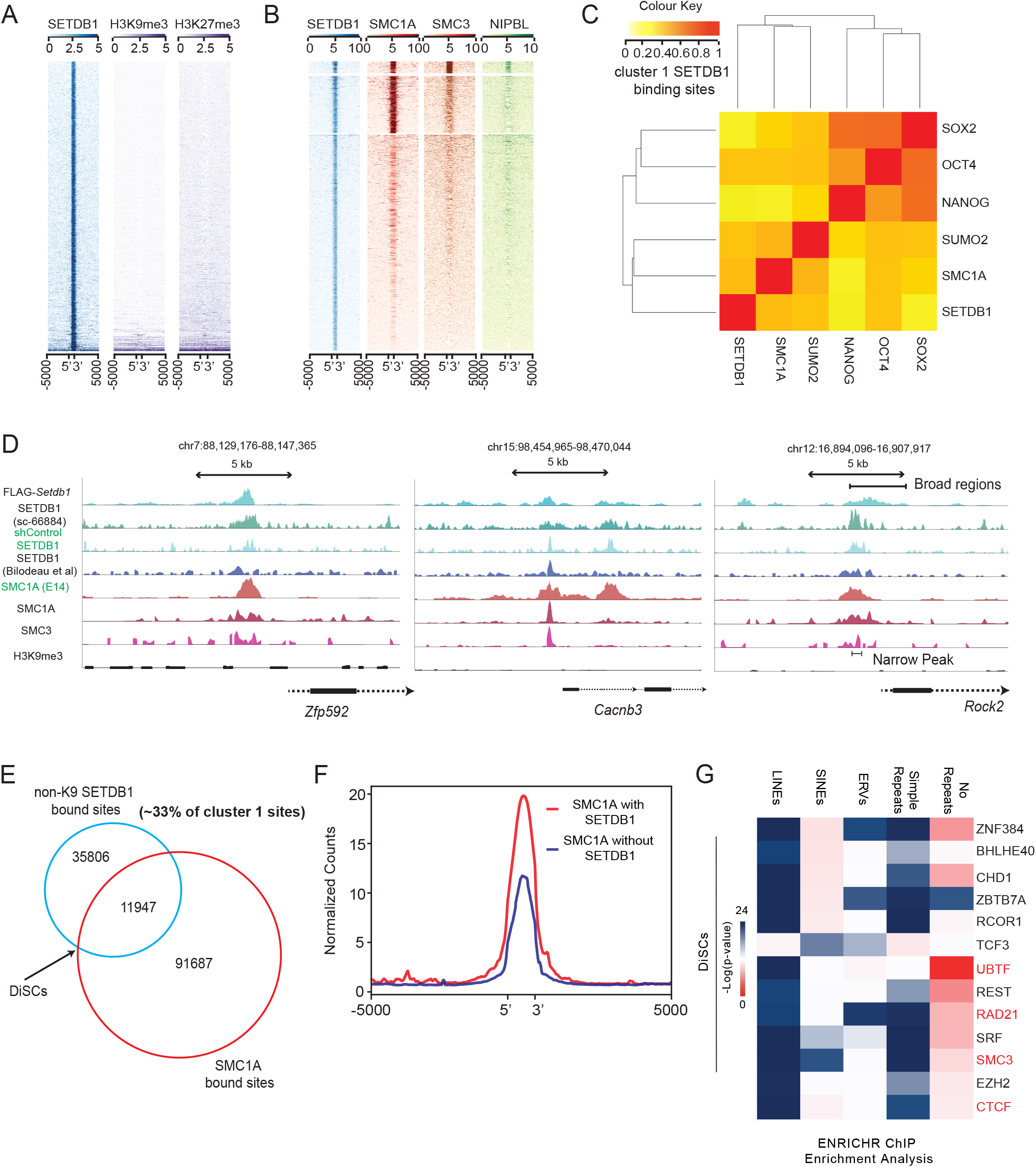
H3K9-independent SETDB1 is enriched at Cohesin-bound sites. **A**, Heat map representing non-canonical SETDB1 binding sites free of repressive H3K9me3 and H3K27me3 (annotated as cluster 1). **B**, Heat map indicating co-binding of Cohesin complex components – SMC1A, SMC3 and Cohesin loading factor – NIPBL on cluster 1 sites. **C**, Clustering analysis for cluster 1 sites bound by the indicated factors (Jaccard coefficient analysis). The colour intensity signifies the strength of the correlation. Red indicates strong correlation, orange indicates medium and significant correlation and yellow represents weak correlation. **D**, UCSC browser view of DiSCs indicating SETDB1 and SMC1A binding profiles (with the use of multiple public datasets as well as replicate ChIP-Seq experiments), with absence of H3K9me3. The left panel represents the presence of broad binding regions for both SETDB1 and SMC1A at DiSCs, contrasted with conventionally detected narrow peaks. **E**, Venn diagram for enhanced DiSC numbers based on broad peak calling for SETDB1 and SMC1A. **F**, Average binding profiles indicating higher levels of SMC1A binding at regions bound by SETDB1 in comparison to genomic regions without SETDB1. **G**, ENRICHR ChIP enrichment analysis for delineation of factors bound to DiSC proximal genes, classified into repeat regions (LINEs, SINEs, ERVs, Simple repeats) and non-repeat regions.

To understand the molecular specifications of cluster 1 SETDB1-bound sites, we performed DNA-binding motif analysis on these sites and identified CTCF-binding motif as the most significantly enriched motif, which were found in ~11% of cluster 1 SETDB1-bound sites (**Supplementary Figure S1D**). Since CTCF regulates genome topology along with the Cohesin complex^63,64^, we next examined whether the Cohesin subunit SMC1A binds to cluster 1 SETDB1 binding sites and found that a total of 2644 such sites (~30% of cluster 1 sites) were highly enriched with SMC1A binding (**Supplementary Table S2**) (**Supplementary Figure S1E**). A similar binding profile was also observed for the Cohesin ring subunit SMC3 and its loading protein NIPBL (**Figure 1B**).

A similarity coefficient-based analysis^65^ between several factors across cluster 1 SETDB1-bound sites revealed that SETDB1 and SMC1A were grouped together at these sites (**Figure 1C**). Visually the binding of SETDB1 and SMC1A at these sites was highly co-localized with overlapping peaks covering similar sized stretches over the genome (**Figure 1D**). As a result of this co-localisation, these sites were annotated as Domains involving SETDB1 and Cohesin (DiSCs). Further visualisation of the DiSCs revealed that some of the sites exhibited broad binding profiles for SETDB1 and SMC1A (**Figure 1D, Supplementary Figure S1F**). To statistically profile these broad regions, a modified peak-calling paradigm for the ChIP-Seq libraries allowed the identification of all broad peaks at the DiSC sites and expanded the list of such sites to 11947 loci (**Figure 1E**) (**Supplementary Table S3**). It is known that histone modifiers such as SETDB1 and topological modifiers such as Cohesin translocate across the genome and hence the broad peak profiles are indicative of an insight into their dynamic locomotion across the genome. Next, a broad versus narrow peak analysis was performed for SETDB1 and Cohesin complex sub-units RAD21 and SMC1A (**Supplementary Figure S1G**). In both cases a substantial percentage of sites were observed to have broad profiles. In contrast, CTCF demonstrated a very high overlap between the two forms of peaks. This suggests that CTCF binding is highly localised, without the characteristic translocating profile displayed by both SETDB1 and SMC1A. Strikingly, SMC1A binding at SETDB1-bound sites was also seen to be at least 2-fold higher as compared to sites without SETDB1 across the entire genome (**Figure 1F**). This was the first indication suggesting the strong molecular association between these two proteins. A detailed dissection of the cluster 1 H3K9me3-free, SMC1A-bound loci i.e. the DiSCs further revealed ~73% of the sites to be non-repeat regions that were distinct from the usual repeats that SETDB1 populates (**Supplementary Figure S1H**). Furthermore, motif analysis of the genes proximal to this non-repeat cluster of DiSCs revealed a high prevalence of CTCF and Cohesin sub-units RAD21, SMC3 as compared to other regions (**Figure 1G**). Other enriched motifs like UBTF at these loci indicate these DiSC-proximal genes to be essential components of embryogenesis and homeostasis^66,67^.

Additionally, we were able to validate our DiSCs by displaying binding of SETDB1 across canonical H3K9me3-bound loci (**Supplementary Figure S1I**) and DiSCs (**Supplementary Figure S1J, Supplementary Figure S1K**) in two public SETDB1 ChIP-Seq datasets^11,32^. An overall correlation of our shControl E14 SETDB1 ChIP-Seq with individual public datasets revealed a significant correlation (spearman~0.50) (**Supplementary Figure S1L**), whereas the overall correlation between the individual public datasets was significantly poorer (**Supplementary Figure S1M**). The lower correlation is expected as a result of the dynamic and transient binding of SETDB1 to the genome – as a result, varying between sample to sample. We were also able to show conserved binding profiles for SETDB1 and SMC1A across public datasets and for different replicate libraries prepared by using multiple antibodies, cell lines etc. (**Figure 1D**, **Supplementary Figure S1F**), thus imparting reproducibility to our findings.

### DiSCs exhibit the presence of a putative SETDB1-Cohesin complex

The binding trends of both SETDB1 and Cohesin (represented by its ring sub-unit SMC1A) were validated by ChIP-qPCR (**Figure 2A, Figure 2B, Supplementary Figure S2A**). We further validated the SETDB1 binding at DiSCs by engineering a FLAG-*Setdb1* cell line (**Supplementary Figure S2B**). ChIP-qPCR validation experiments using the FLAG-*Setdb1* cell line showed consistent SETDB1 binding at the DiSC sites (**Figure 2C and Supplementary Figure S2C**). Additional evidence for SETDB1 and SMC1A co-binding at these sites was provided by sequential ChIP experiments, with no signal seen at loci that were only SETDB1-bound (*Polrmt*) or SMC1A-bound (*Vav2*) (**Figure 2D**). SETDB1 and SMC1A were also seen to strongly interact with each other in over-expression Co-IP experiments performed in HEK293T cells (**Figure 2E**), thereby delineating the presence of a potential regulatory complex. Furthermore, IP-MS datasets for endogenous E14 cells for both SMC1A and SETDB1 revealed the presence of SETDB1/SMC1A/SMC3 in both IP-MS datasets (**Figure 2E**)(**Supplementary Table S4**). Ablation of *Setdb1* using an shRNA construct, showed substantial reduction in SETDB1 binding at the DiSCs, indicating that the binding was specific (**Supplementary Figure S2D, Supplementary Figure S2E**). ChIP experiments performed after *Smc1a* knock-down (**Supplementary Figure S2D, Supplementary Figure S2F**) also confirmed the binding specificity of Cohesin at DiSCs.

**Figure 2.**
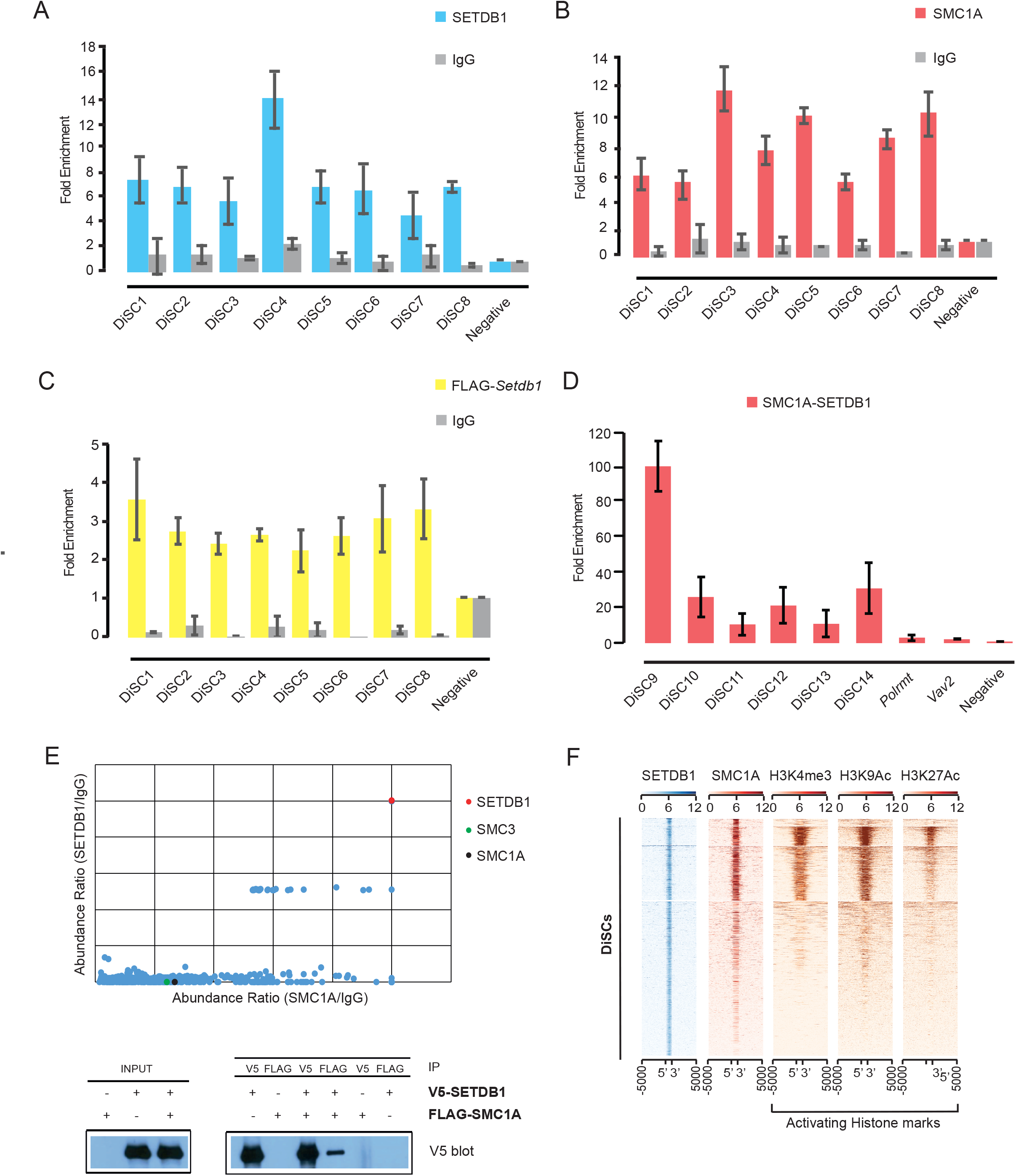
SETDB1 and SMC1A strongly co-exist at DiSCs. **A**, ChIP-qPCR analysis of SETDB1 binding at DiSCs (n=3, Error bars represent standard deviation). **B**, ChIP-qPCR analysis of SMC1A binding at DiSCs (n=3, Error bars represent standard deviation). **C**, ChIP-qPCR analysis of SETDB1 binding at DiSCs as seen by using the FLAG-*Setdb1* cell line (n=3, Error bars represent standard deviation). **D**, Sequential ChIP-qPCR indicating the co-binding of SMC1A and SETDB1 at DiSCs. *Polrmt* and *Vav2* indicate SETDB1 and SMC1A only binding sites, thereby serving as sequential ChIP controls (n=3, Error bars represent standard deviation). **E**, (Top) IP-MS datasets indicating the abundance ratios compared to IgG for SETDB1 (y-axis) and SMC1A IP (x-axis) in mESCs (E14) indicating the presence of SETDB1, SMC1A, SMC3 in both datasets. (Bottom) Western blot for V5-SETDB1 following overexpression of V5-SETDB1 and FLAG-SMC1A in the HEK293T cells followed by co-immunoprecipitation indicating the strong interactions between SETDB1 and SMC1A. **F**, Heat maps depicting the binding profiles of active histone marks (H3K4me3, H3K9Ac and H3K27Ac) at the DiSCs.

As further validation of the importance of these sites in pluripotent cell types, we carried out reprogramming of mEF (mouse embryonic fibroblasts) cells to induced pluripotent stem cells (iPSCs)^68^ and confirmed SETDB1 and SMC1A binding at these sites in the reprogrammed cells (**Supplementary Figure S2G**). This was a clear indication towards the potentially conserved nature of these sites in pluripotent cell types. *Setdb1* and *Smc1a* were also seen to co-express significantly across varied tissue and cell types in the ENCODE dataset^39^ (r=0.75) (**Supplementary Figure S2H**) and moreover, the DiSCs were also seen to be moderately conserved across a set of 30 vertebrate species as per their PhastCon scores (**Supplementary Figure S2I**). These findings successfully demonstrate an unprecedented but strong association between SETDB1 and Cohesin at the DiSCs.

Since the DiSCs were bereft of repressive histone marks – which are common associates of SETDB1, we profiled other activating histone marks across the DiSC sites, and strikingly observed a presence of H3K4me3 and H3K27Ac, H3K9Ac (which are marks enriched at promoters and enhancers) (**Figure 2F**, **Supplementary Figure S2J**). This was a strong indication of the SETDB1-Cohesin complex at DiSCs being involved in transcriptional regulation.

### DiSCs mediate transcriptional control of their adjacent genes

Following on from the enrichment of activation histone marks proximal to DiSCs, we were curious about gene proximity to the DiSCs. Therefore, a profile was constructed based on their distance from the closest transcription start sites (TSS) (**Figure 3A**). Majority (51%) of the DiSCs were seen to be distal with respect to TSS i.e., at 10000 bp or more, whereas around 28% can be found within 1000 bp of the nearest TSS. Hyper-geometric tests confirmed that most of the DiSCs were located proximal to the promoters (~33%) and 5’UTR (~6%), whereas ~8% were located downstream (1000-3000 bp) to the genes (**Supplementary Figure S3A**). Since significant numbers of DiSCs were proximal to the gene regulatory regions, we postulated that they were likely to engage in transcriptional activity. DiSC sites were selected and successfully tested for transcriptional activity using a luciferase assay (**Supplementary Figure S3B**).

**Figure 3.**
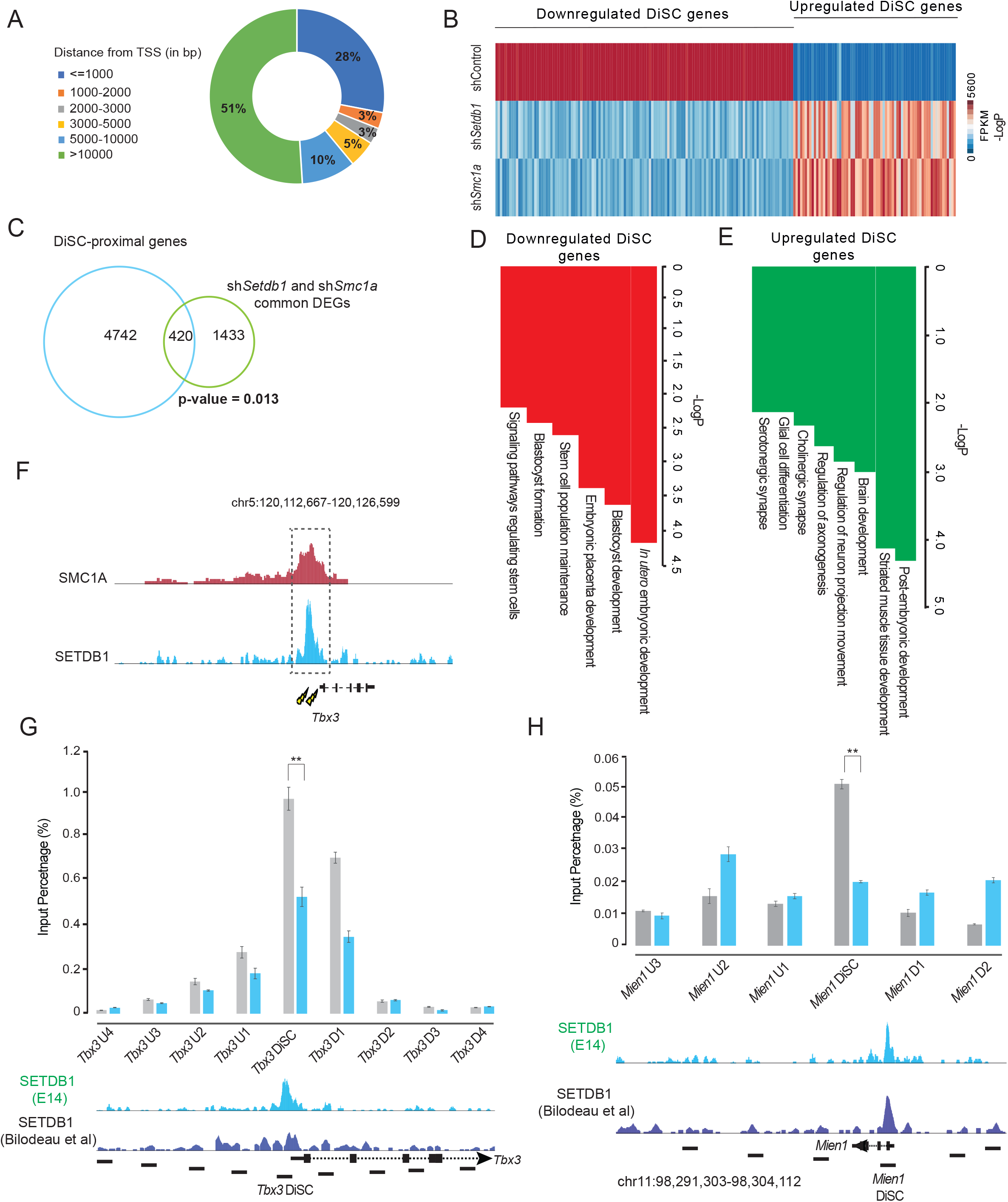
SETDB1 and SMC1A at DiSCs co-regulate transcriptional profiles. **A,** Distribution of DiSCs based on their distance to the nearest transcription start site (TSS) delineating either a promoter proximal or distal profile for these sites. **B**, Heat map delineating the DEGs that are seen to be commonly dysregulated by SETDB1 and SMC1A, at DiSCs. **C**, Venn diagram representing the overlap of commonly dysregulated genes for SETDB1 and SMC1A with DiSC-proximal genes – revealing 420 such statistically significant genes. **D**, Gene ontology (GO) analysis for downregulated DiSC-proximal genes. **E**, Gene ontology (GO) analysis for upregulated DiSC-proximal genes. **F**, Schematic visualizing the CRISPR-Cas9 based targeting of SETDB1-SMC1A bound sites at DiSCs, proximal to the *Tbx3* gene. **G**, Locally depleted SETDB1 enrichment at targeted *Tbx3* DiSC site, with little or no impact on SETDB1 binding around the target site (n=3, Error bars represent standard deviation). Twotailed t-test was used for statistical analysis. [p-value < 0.05, **p-value < 0.01]. **G**, Locally depleted SETDB1 enrichment at targeted *Mien1* DiSC site, with little or no impact on SETDB1 binding around the target site (n=3, Error bars represent standard deviation). Two-tailed t-test was used for statistical analysis. [p-value < 0.05, **p-value < 0.01].

To further explore gene regulatory roles of the DiSCs, we prepared RNA-Seq libraries for shControl, sh*Setdb1* and sh*Smc1a* cells. These libraries showed depleted expression of the shRNA-targeted genes (**Supplementary Figure S3C**). In the *Setdb1*-depleted libraries, 892 genes (proximal to the DiSCs) were upregulated while 1322 genes were downregulated. In the *Smc1a*-depleted libraries, 998 genes (proximal to the DiSCs) were upregulated and 1338 were downregulated (**Supplementary Figure S3D**). While a high number of differentially expressed genes (DEGs) i.e. 800, in the sh*Setdb1* RNA-Seq were seen to be DiSC-proximal genes (**Supplementary Figure S3E**), the majority of genes proximal to SETDB1, H3K9me3 co-bound loci were upregulated rather than downregulated as expected (**Supplementary Figure S3F, Supplementary Figure S3G**). A substantial percentage of the dysregulated genes were seen to be impacted by both *Setdb1* and *Smc1a* depletion i.e. 420 of the common dysregulated genes overlapped with DiSC-proximal genes (**Figure 3C**). Out of this number 264 genes were downregulated and 156 of the upregulated genes were common between the sh*Setdb1* and sh*Smc1a* libraries (**Supplementary Figure S3H**) (**Supplementary Table S5**) On the other hand genes commonly dysregulated between sh*Setdb1* and sh*Smc1a* did not overlap significantly with SETDB1, H3K9me3 co-bound genes (**Supplementary Figure S3I**). Gene ontology (GO) analysis for the commonly downregulated DiSC-proximal genes revealed enrichment for core stem cell maintenance and survival processes – blastocyst development, stem cell population maintenance, and broadly signalling pathways regulating stem cells (**Figure 3D**). Similar analysis for the upregulated DiSC-proximal genes displayed functions relevant to post-embryonic development, brain development, glial cell differentiation (**Figure 3E**). GO for the SETDB1-H3K9me3 regulated genes was seen to be distinctly different as it was enriched with varied metabolic processes, cell maintenance functions and other differentiation related processes (**Supplementary Figure S3J**). The GO analysis at the DiSCs was strongly supported by the factors enriched at the DiSC-proximal upregulated genes – TAF1, ATF2, MYC, all of which have been implicated in distinct roles in neuronal development and NPC differentiation^69,70^ (**Supplementary Figure S3K**).

In order to show a direct correlation between DiSCs and their target gene expression, two DiSCs proximal to the genes, *Tbx3* (**Figure 3F**) and *Mien1*(**Supplementary Figure S3L**), were targeted for knock-out (KO) using CRISPR-Cas9. After disrupting the SETDB1-binding site upon the targeted DiSC, the expression of the proximal gene was significantly downregulated in accordance with the trends previously observed in the KD RNA-Seq libraries (**Supplementary Figure S3L, Supplementary Figure S3M**). Moreover, the disruption of the proximal DiSC site contributed to gene dysregulation via a localized decline in enrichment of SETDB1 at the targeted site (**Figure 3G, Figure 3H**), thereby indicating DiSCs to be gene expression regulatory elements functioning via the presence of SETDB1 and Cohesin.

### SETDB1 and Cohesin binding are mutually interdependent and critical for genome topology at DiSCs

Having ascertained the interactions between SETDB1 and Cohesin at the DiSCs and their combined impact on proximal gene dysregulation, we next wished to explore the nature of dependency between these two ‘unlikely’ partners. Since SMC1A binding was seen to be stronger at SETDB1-bound genomic regions, we performed a genome-wide analysis of SMC1A binding after depletion of SETDB1. Interestingly, we observed drastic decrease of Cohesin binding at more than half the DiSCs after the knock-down of SETDB1 (**Figure 4A, Supplementary Figure S4A**). At the impacted DiSCs, the decrease in SMC1A enrichment was seen to be at least 2-fold for sites that were regarded as significant (**Figure 4B, Figure 4C**). Some observable decrease in SMC1A levels after SETDB1 KD was observed across other tested sites such as SMC1A-H3K9me3 and SMC1A-CTCF co-bound sites, however the decreased SMC1A enrichment was most pronounced across the DiSCs (**Supplementary Figure S4B, Supplementary Figure S4C**). ChIP-qPCR based validation experiments further showed both Cohesin complex sub-units, SMC1A and RAD21 as having reduced enrichment at the tested DiSCs after SETDB1 KD (**Figure 4D and Supplementary Figure S4D**).

**Figure 4.**
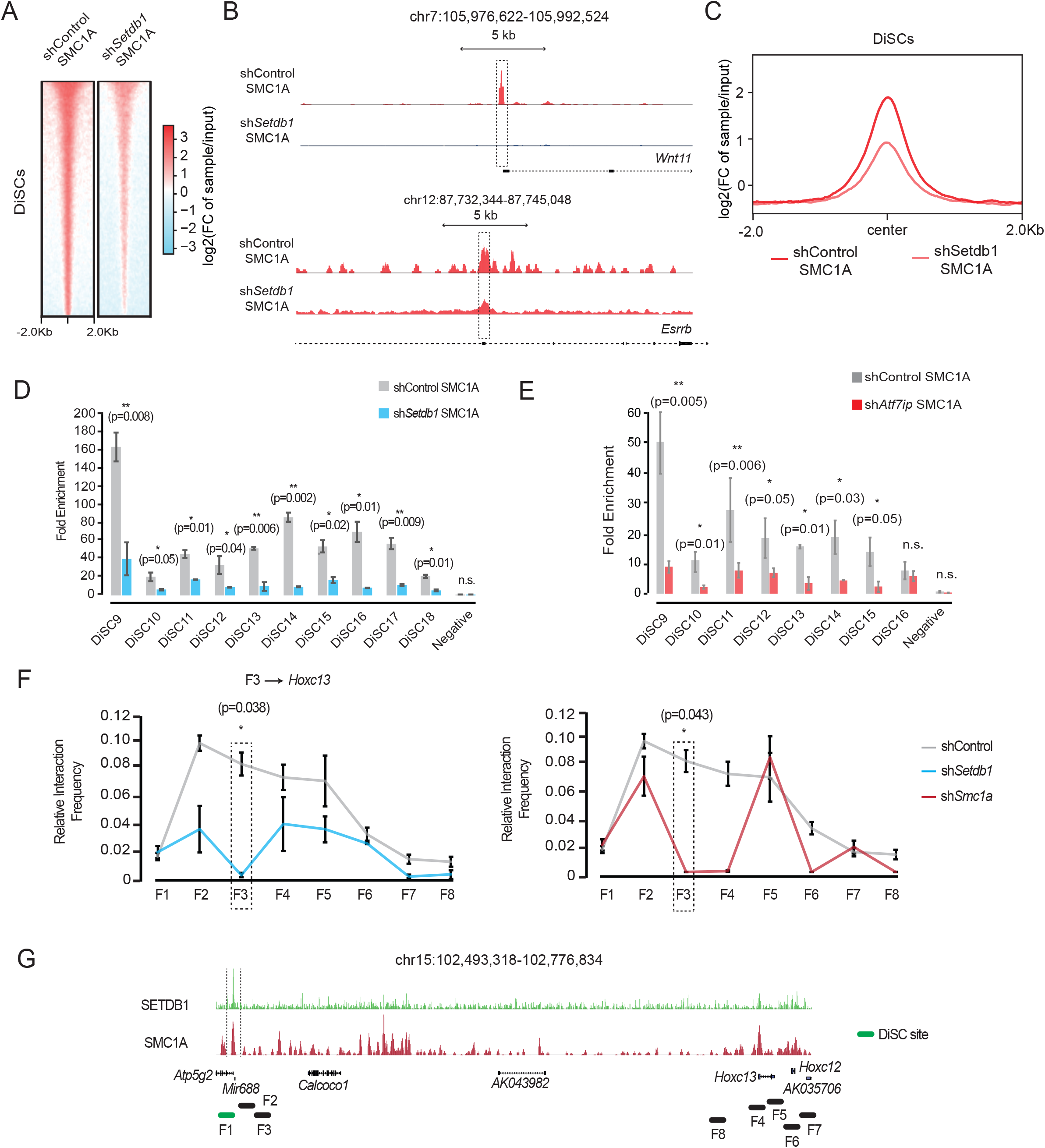
SETDB1 dictates Cohesin levels at the DiSCs. **A**, Heat map delineating the decreased levels of SMC1A upon SETDB1 KD across the DiSCs. **B**, UCSC view of the drastic decline of SMC1A (Cohesin) binding after KD of SETDB1 at the visualised DiSCs. **C**, Average enrichment profile for SMC1A enrichment in shControl and sh*Setdb1* cells representing ~2-fold decrease in Cohesin levels across DiSCs upon SETDB1 KD. **D**, ChIP-qPCR based validation of the decrease in SMC1A binding at the DiSCs after KD of SETDB1 (n=3, Error bars represent standard deviation). Two-tailed t-test was used for statistical analysis. [p-value < 0.05, **p-value < 0.01]. **E**, ChIP-qPCR based validation of the decrease in SMC1A binding at the DiSCs after KD of SETDB1 (n=3, Error bars represent standard deviation). Two-tailed t-test was used for statistical analysis. [p-value < 0.05, **p-value < 0.01]. **F**, 3C-qPCR interaction profiles around *Hoxc13* genomic locus which is located proximal to a DiSC, upon KD of SETDB1 (left) and SMC1A (right). (n=3, Error bars represent standard deviation). Two-tailed t-test was used for statistical analysis. [*p-value < 0.05, **p-value <0.01]. **G**, UCSC view of the DiSC site (proximal to the *Mir688* gene) and 3C-qPCR amplified fragments in the vicinity of the *Hoxc13* gene.

SETDB1 activity is stabilised in cells by the protein ATF7IP – which sequesters SETDB1 and protects it from proteosomal degradation^6,7^. Therefore, SETDB1 and ATF7IP co-exist and function together at a molecular level. To associate this with the SETDB1-SMC1A relationship, we performed KD for ATF7IP, and observed a decline in the enrichment levels of SMC1A and RAD21 at tested DiSC sites (**Figure 4E, Supplementary Figure S4E, Supplementary Figure S4F**). This clearly implied that the ATF7IP-SETDB1-Cohesin cascade existed as a determinant of Cohesin binding across the genome and thereby could be essential for gene regulation and genome topology. Despite the decreased binding of SMC1A across the genome after SETDB1 depletion, there was no significant change in the expression levels of the Cohesin ring sub-unit genes – *Smc1a, Smc1b, Smc3* (**Supplementary Figure S4G**).

Cohesin is specifically enriched at TADs (Topologically Associating Domains) and other topological structures across the genome^71^. Since the enrichment of SMC1A at DiSCs is a function of SETDB1 levels, we next explored whether the presence or absence of SETDB1 and SMC1A were influential in regulating topological interactions at the DiSCs, by performing a 3C-qPCR assay. We identified two DiSC-proximal genes *Hoxc13* and *Ppp1r1b* that were seen to be dysregulated in sh*Setdb1* and sh*Smc1a* RNA-Seq samples, and used them to probe the topological interactions in their vicinity (**Supplementary Figure S4H**). Upon the depletion of SETDB1 or SMC1A, we observed a drastic reduction in the prevalence of genomic interactions (**Figure 4F, Supplementary Figure S4I, Supplementary Figure S4J**). The presence of a proximal DiSC (indicated by SETDB1, SMC1A binding profiles) could be clearly seen for the tested sites, thereby asserting that the changes in localized expression were accompanied by topological changes in the vicinity (**Figure 4G**, **Supplementary Figure 4J**). Hi-C data clearly displayed interactions around these two tested gene loci and showed presence of loops dictated by Cohesin (as per public Cohesin Hi-ChIP data^52^) (**Supplementary Figure S4K, Supplementary Figure S4L**). These findings cumulatively indicated that the SETDB1-Cohesin partnership was crucial for dictating genome topology at the DiSCs, and thereby was also critical for gene expression regulation.

### DiSCs comprise critical topological structures in the genome dictated by SETDB1

Overlaying DiSCs upon Cohesin Hi-ChIP data for the E14 mouse ES cell line implicated the DiSCs in TAD and loop architecture (**Figure 5A, Supplementary Figure S5A)**. To further probe the roles of SETDB1 and Cohesin at the DiSCs, we prepared Hi-C libraries after KD of *Setdb1* and *Smc1a*, with the shControl library as a wild-type control. Strikingly, it was seen that the depletion of SETDB1 impacted TADs across the genome adversely, while expectedly SMC1A (Cohesin) loss depleted topological domains drastically (**Supplementary Figure S5B**). Both SETDB1 and SMC1A loss also depleted topological interactions at DNA loops and adversely impacted the loops located at DiSCs (**Figure 5B, Supplementary Figure S5C**) (**Supplementary Table S6**).

**Figure 5.**
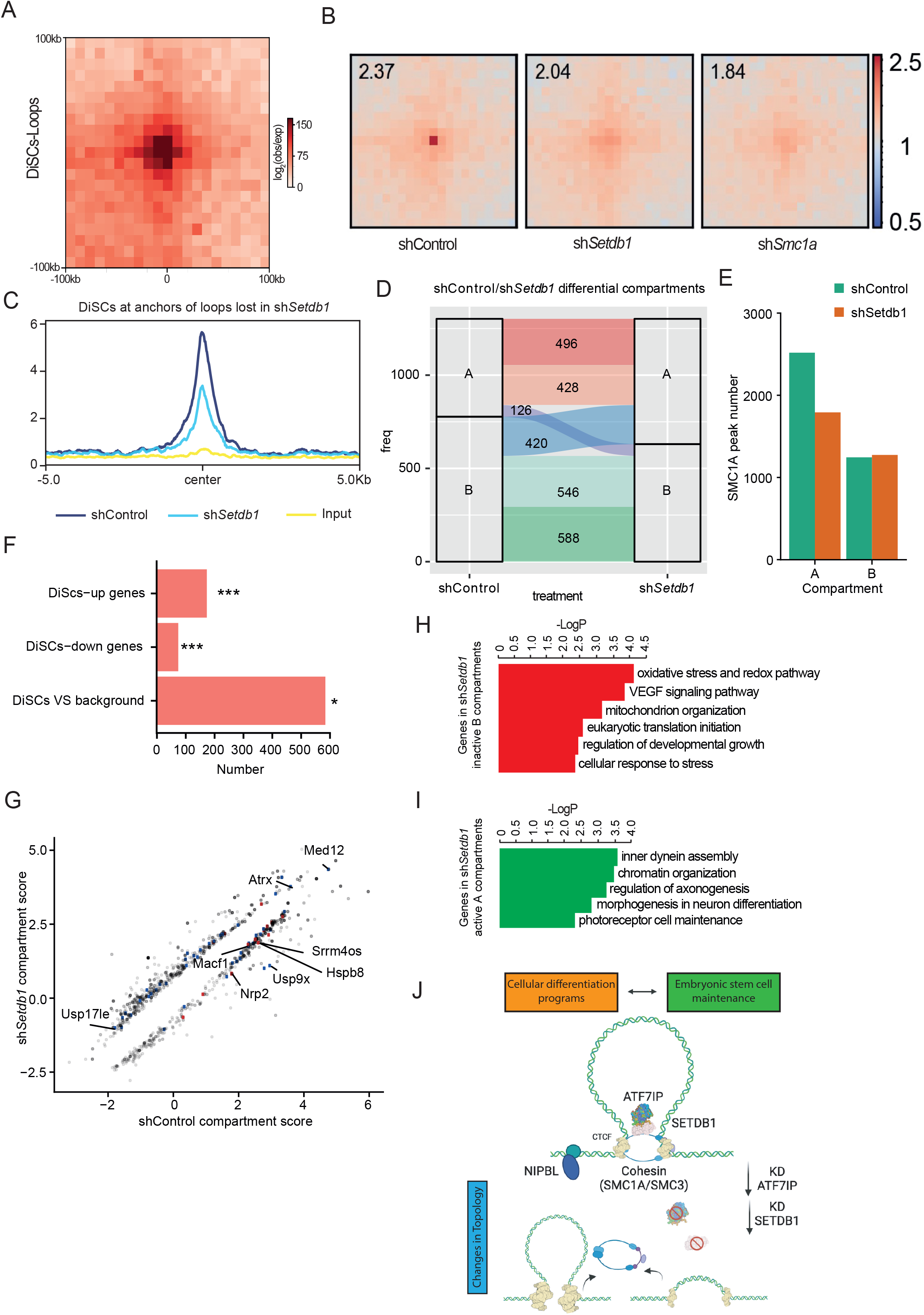
SETDB1-Cohesin constitute a topological axis that regulates gene expression. **A,** Overlap of significant loops detected in Cohesin HiChIP with DiSC sites at a 10kb resolution. **B,** Depletion of topological interactions at DiSCs upon KD of SETDB1 and SMC1A. **C,** Average enrichment profiles for SMC1A binding in shControl and sh*Setdb1* cells across loop anchors that show depleted topological profiles upon KD of SETDB1. **D,** Transition plot delineating compartment switches (A-B, B-A) between shControl and sh*Setdb1* cell states. 126 A-B compartment switches and 420 B-A compartment switches were seen to occur. **E,** Overlap of SMC1A peaks with A and B compartments classified in the shControl and sh*Setdb1* samples. **F,** Enrichment of DiSCs over an equal number of genomic sites used as background at loop anchors in shControl Hi-C datasets. Plot also represents significant enrichment of upregulated and downregulated DiSC genes proximal to loop anchors. **G,** Scatter plot delineating the presence of upregulated <u>(blue)</u> and downregulated <u>(red)</u> genes in differential compartments that arise as a result of sh*Setdb1* treatment, contrasted with shControl Hi-C datasets. **H,** Gene ontology (GO) analysis for downregulated genes seen in differential compartments arising as a result of sh*Setdb1* treatment. **I,** Gene ontology (GO) analysis for upregulated genes seen in differential compartments arising as a result of sh*Setdb1* treatment. **J,** Schematic representation of bi-modal regulation of the genome by SETDB1. This non-canonical model depicts SETDB1 (stabilised by ATF7IP) associating with the Cohesin complex at topological structures (DNA loops, TADs) across the genome and exerting control over them by regulating Cohesin binding to DNA (right). In totality, these two modes of SETDB1 activity control genome topology and gene expression, striking a fine balance in the ES state between embryonic stem cell maintenance and differentiation to other lineages (neuronal, muscle etc.).

The loop anchors that are located at DiSCs concurrently show depletion of SMC1A enrichment after SETDB1 depletion, thereby connecting the loss of topology at DiSCs with the SMC1A depletion upon SETDB1 KD (**Figure 5C**). The distinct topological impact of SETDB1 and SMC1A depletion was visible upon studying the Directionality Index (DI) profiles of shControl, sh*Setdb1* and sh*Smc1a* libraries – thereby indicating genome-wide topology changes, inclusive of the DiSCs (**Supplementary Figure S5D**).

DiSC-proximal genes were seen to be upregulated as well as downregulated upon depletion of SETDB1 and SMC1A. Typically, SETDB1 depletion is known to cause widespread upregulation of genes as a result of the ablation of its deposited H3K9me3 mark. However, the duality of gene expression at DiSCs via SETDB1 and Cohesin was a unique defining trait for the DiSCs. In order to study this further, we profiled the active (A) and inactive (B) topological compartment profiles in our Hi-C datasets, and observed A-B and B-A compartment switches between shControl and sh*Setdb1* Hi-C datasets (**Figure 5D, Supplementary Figure S5E**). As a comparison, sh*Smc1a* Hi-C data also demonstrated widespread compartment switches, as has been known to occur during Cohesin depletion^72,73^ (**Supplementary Figure S5F**).

We observed comparable numbers of Cohesin-bound sites enriched in the inactive B compartments of both shControl and sh*Setdb1* Hi-C datasets, whereas there was a small drop (2500 to 1800) in the number of Cohesin peaks enriched at the active A compartments in sh*Setdb1* as compared to the shControl (**Figure 5E**). This drop was indicative of decreased active compartments comprising Cohesin in the sh*Setdb1* cells, consistent with the loss of Cohesin seen in sh*Setdb1* cells. Upon comparing DiSCs overlapping with loop anchors versus background genomic regions, we observed a significant enrichment for the former set. Additionally, we also observed the presence of both upregulated and downregulated genes associated with DiSCs at the loop anchors, thereby justifying the duality of gene expression control via SETDB1 at the DiSCs. These DiSC-associated genes and DiSCs were much more significantly enriched at loops as compared to SETDB1-H3K9me3 co-bound regions (**Figure 5F**). We could also see the co-operative influence of SETDB1 and Cohesin at the DiSCs, at a number of dysregulated genes found in differential topological compartments upon KD. For instance, the gene *Ppp1r1b*, which is downregulated upon sh*Setdb1* and sh*Smc1a* treatment, was also representative of topological interactions that were lost in its vicinity upon KD of SETDB1 and SMC1A. Similarly, the gene *Pim2*, which was upregulated, displayed interactions arising in its vicinity upon KD of SETDB1 and SMC1A (**Supplementary Figure S5G**).

In line with our hypothesis for the SETDB1-Cohesin axis as a core regulator of genome topology and transcriptional regulation, we found DiSC-associated genes - which were upregulated and downregulated, to be associated with differential compartments in the sh*Setdb1* Hi-C dataset (**Figure 5G**) (**Supplementary Table S7**). The genes that were downregulated in the sh*Setdb1* compartments were seen to be engaged in functions of cell maintenance, stress response and growth (**Figure 5H**), while the upregulated genes were central to neuronal development and chromatin organisation (**Figure 5I**). These GO trends agreed with the wider functional trends seen earlier for DiSC-proximal genes that were dysregulated, with upregulated genes playing key roles in stem cell differentiation to neurons, muscle etc., whereas the downregulated genes were critical for stem cell metabolism and maintenance.

## Discussion

We have identified through our extensive analysis of SETDB1 ChIP-Seq data - novel regions known as DiSCs which are bound by SETDB1 and Cohesin. These unique regions were seen to be proximal to genes and bereft of all known repressive histone marks, implying their significantly different functions with respect to SETDB1 and its known roles. The strong cooccupancy of SETDB1 and Cohesin at these sites suggested their inter-dependency and indeed it was strikingly seen that Cohesin enrichment at these sites was heavily reliant on SETDB1 binding. Furthermore, depletion of ATF7IP, which is an upstream regulator and stabiliser for SETDB1, also cascaded on to depletion of SMC1A and the Cohesin complex at large. Therefore, we postulated the existence of a SETDB1-Cohesin regulatory system, wherein SETDB1 functioned as an upstream regulator of Cohesin at the DiSCs. This hypothesis was further strengthened by the impact on DiSC-proximal gene expression as well as localized genome topology after SETDB1 depletion in the cells, which are effects of the decline in localized Cohesin levels. KD SETDB1 Hi-C data revealed the functional significance of this SETDB1-Cohesin regulatory axis by establishing DiSCs as significant topological structures that are impacted upon SETDB1 loss, leading to gene expression regulation. Overall, we have been able to profile a unique regulatory relationship between SETDB1 and Cohesin, thereby leading to the understanding that SETDB1 is a critical regulator of cell fates and cell lineage, not just via its deposited repressive H3K9me3 marks, but also by virtue of exerting control on Cohesin association to the genome. By combining with Cohesin, SETDB1 establishes the DiSCs as unique topological and functional domains that are crucial on the one hand in maintaining the stem cell state, but can also serve as triggers for differentiation and cellular transformation to alternative lineages (neurons, muscles etc.).

SETDB1 has not been extensively studied, outside of its conventional histone methyltransferase function. A recent report indicated that SETDB1 could be located proximally to a large neuron-specific topological domain and be potentially involved in its regulation^15^, however this too was linked to its H3K9me3-dependent role. The unique nature of the DiSCs stems from the fact that they are free of SETDB1-associated repressive histone marks such as H3K9me3 and H3K27me3, and are rather co-occupied by Cohesin. The co-existence of SETDB1 and SMC1A at these domains is extensive and highly functional, suggesting that they are acting together as a complex.

Many of the DiSCs have unique broad binding profiles which we suspect represents the dynamic binding of SETDB1 and Cohesin to chromatin. Cohesin complex is known to slide and translocate across the length of chromatin^74–76^. As such, they typically exhibit broader binding profiles. We have been able to validate this observation for RAD21 – another Cohesin sub-unit, whereas this does not seem to be the case for a transcriptional regulator such as CTCF, which binds to a specific motif. It is feasible that the broad binding profile of SETDB1, as an enzyme, might be a consequence of its transient and rapidly changing association to its substrate proteins that are bound across the genome. Remarkably, the broad binding profile of SETDB1 was reproducible across different antibody reagents and a FLAG-*Setdb1* ES cell line we had engineered. For Cohesin, in accordance to the model of loop extrusion, the broad peak profiles could also arise from the transitional nature of the CTCF-Cohesin associations^77^. DiSCs are enriched with activating histone marks such as H3K4me3 and hence comprise of transcriptionally active regions, unlike the regular association of SETDB1 with repressive marks^78^. Therefore, our discovery of the DiSCs demarcates them as unique, functionally relevant regions.

Recent studies have suggested that Cohesin has a prominent role in terms of regulating the 3D genome^29^, and have also implicated genome topology in tissue-specific transcriptional regulatory roles^79^. Our analysis of Hi-C, KD Hi-C and HiChIP datasets implied that the DiSCs are indeed topological structures that regulate chromatin architecture. Interestingly, a significant proportion of the DiSCs with disrupted Cohesin binding after SETDB1 ablation were accompanied by a loss of genome topology, subsequently leading on to proximal gene dysregulation. Out of these dysregulated genes a substantial percentage were commonly impacted by the ablation of SETDB1 and SMC1A. The functions of the commonly downregulated functional DiSC genes were seen to be related to stem cell maintenance and embryonic development as well as other metabolic and cellular processes such as RNA and DNA metabolism, translation. This aligns with previous findings which suggested that SETDB1 was essential in the maintenance of pluripotency^33^. Previous findings discussed SETDB1-regulated repression of lineage-specific or developmental genes by deposition of the H3K9me3 mark. The regulation of pluripotency specific genes, pathways and processes via topological domains such as DiSCs adds a new dimension to its role in the regulation chromatin architecture. Upregulated DiSC proximal genes, on the other hand had lineage specific functions such as muscle and neuronal lineage development. On a functional basis, the discovery of DiSCs in this study, has thus paved a new mechanistic and topological aspect to gene expression regulation for differentiated cell types (such as neurons) as well as for the maintenance of pluripotency by SETDB1 through its control over Cohesin binding at these domains. The topological link of SETDB1 at the DiSCs was further strengthened by observing a strong loss of topological interactions at DiSC loop anchors upon ablation of SETDB1. The topological impact exerted by SETDB1 via Cohesin also contributed to compartment switches and gene dysregulation as a result.

Together, DiSCs exhibit a non-canonical model for the binding of SETDB1 (**Figure 5J**) on the genome, without its classical association to repressive histone marks. The regulation of Cohesin binding by SETDB1 at these sites as well as the subsequent impact on gene dysregulation are tightly linked to the local genomic 3D topology and architecture^80^. DiSCs also possess lineage specific gene regulation that are chiefly controlled by SETDB1 and Cohesin. Recent studies have shown that histone-specific methyltransferases have been found to methylate non-histone proteins in some instances^81,82^. SETDB1 too, has recently been shown to act on P53 as well as AKT as a non-histone protein methyltrasnferase^18,8,9^. We also speculate that the role of SETDB1 in controlling Cohesin binding at the DiSCs is related to its enzymatic role. To this end, we were also able to profile methylated lysine residues on Cohesin sub-units SMC1A, SMC3 (data not shown). Most methylated lysine and arginine residues on proteins exhibit biological relevance in context of their functional and structural stability^83–85^. Drawing inferences from the clear correlation between the presence of SETDB1 at the DiSCs, and its influence on the binding of Cohesin, there remains a mechanistic possibility of SETDB1 influencing Cohesin via its methyltransferase activity, which is further influenced by its complex with ATF7IP. However, this biochemical validation of the SETDB1-Cohesin axis needs further experimental validation and probing, and was not within the current functional and topological scope of our exploration of the DiSCs.

Therefore, this study reports the discovery of DiSCs, revealing these previously undescribed domains of the genome that act as localized transcriptional factories, regulating specific lineages and functions via topological structures cumulatively controlled by SETDB1 and Cohesin. The impact of DiSC perturbation on gene expression linked to neural and muscle system development is particularly fascinating since it provides an avenue to explore facilitation of differentiation processes by targeting of specific DiSCs. These findings also illustrate how regions like DiSCs could prove to be critical regulators of cell fate and lineage. The broad binding profiles of Cohesin and SETDB1 identified from this study, suggests that transient and dynamic chromatin binders may exhibit such a hallmark in its binding to the genome. Our study also delineates the versatility of SETDB1 beyond its traditional repressive role, suggesting that it dictates chromatin topology and exerts gene expression control across the genome independent of its histone H3K9-methylation capabilities.

## Data Availability

All the next-generation sequencing data have been submitted to GEO under the accession GSE123245. All other data can be made available by the corresponding authors upon reasonable request.

## Acknowledgements

We sincerely thank Dr. Fang Haitong for technical assistance in the study. We also thank Dr. Hao Fei Wang, Dr. Qiaorui Xing and Dr. Nareshwaran Gnanasegaran and Aloysius Quek for their helpful suggestions as well as technical assistance. We also thank Bobby Tan from the Genome Institute of Singapore for providing the SETDB1 antibody.

## Author Contributions

T.W., C.E.F. designed and performed the research, analyzed the data and wrote the paper. Y.Y.Z., B.S.Q.H., D.L.X., Q.B., Z.H.Z., X.B., K.Y.S.T. designed and conducted research. H. H.N., D.S.T.O., J.J.H.C., A.S., M.J.F., J.C., H.L. and J.X. analyzed data. Y-H.L. designed research, analyzed data and wrote the paper.

## Supplementary Datasets

**Supplementary Table S1:** shRNA oligos used for KD of *Setdb1, Smc1a, Atf7ip*

**Supplementary Table S2:** cluster 1 SETDB1 binding sites (free of H3K9me3) overlapping with Cohesin

**Supplementary Table S3:** Combined list of DiSC regions with gene annotations (list of expanded DiSCs after broad peak calling)

**Supplementary Table S4:** PPIs and protein partners detected for IP-Mass Spectrometry assays for the endogenous SETDB1 and SMC1A proteins in the mESC (E14) cell line.

**Supplementary Table S5:** List of genes dysregulated by SETDB1 and Cohesin proximal to DiSCs

**Supplementary Table S6:** Differential interactions observed between shControl, sh*Setdb1* and sh*Smc1a* Hi-C libraries

**Supplementary Table S7:** Differential compartments (A and B) observed between shControl and sh*Setdb1* Hi-C libraries along with gene annotations

