## Supplemental Information for "DiSCs – Domains involving SETDB1 and Cohesin are critical regulators of genome topology and stem cell fate"

Supplementary Figure 1

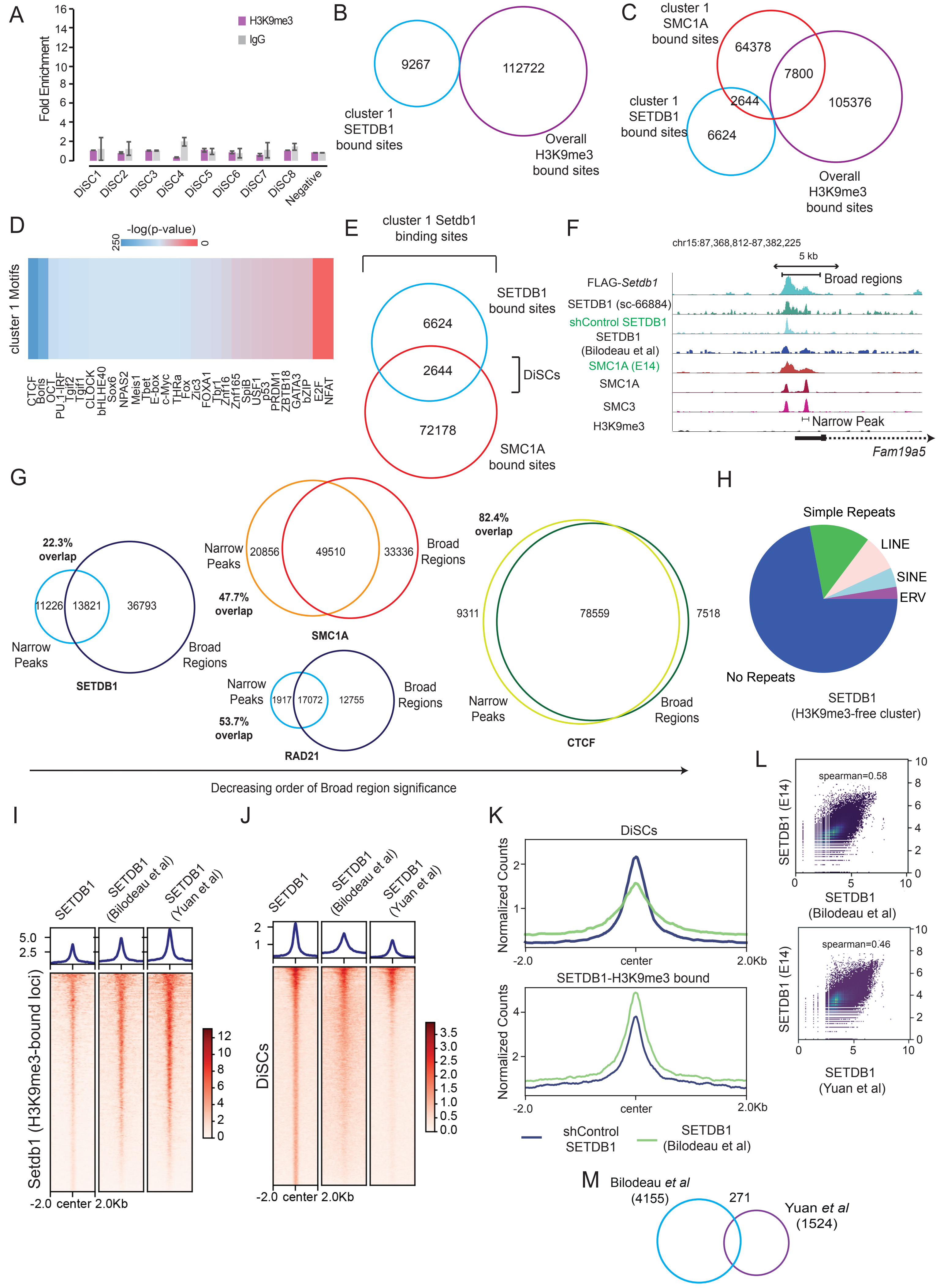

**Supplementary Fig.1 | SETDB1 binds to DiSCs with Cohesin and without H3K9me3 reproducibly.** **A**, ChIP-qPCR validation of the lack of H3K9me3 binding at DiSCs. (n=3, Error bars represent standard deviation). **B**, Overlap of cluster 1 SETDB1-bound sites with repressive mark H3K9me3-bound sites. **C**, Overlap of cluster 1 SETDB1-bound and SMC1A-bound sites with repressive mark H3K9me3-bound sites. **D**, Motifs enriched at cluster 1 SETDB1 binding sites as identified by HOMER, with CTCF being the top-enriched motif. **E**, Overlap of cluster 1 SETDB1-bound sites with SMC1A-bound sites, representing the DiSCs, prior to broad peak-calling for both SETDB1 and SMC1A. **F**, UCSC screenshots representing different SETDB1 libraries (from top to bottom: SETDB1 (FLAG-*Setdb1* cell line), SETDB1 (sc-66884), SETDB1 (shControl library), SETDB1 (Bilodeau *et al.*, 2009), showing binding across DiSC sites. Binding profiles of Cohesin sub-units SMC1A and SMC3 have also been represented. Altogether, broad and narrow binding profiles can be contrasted across these datasets. **G**, Overlap of narrow-style and broad-style peaks for SETDB1 (left), SMC1A and RAD1 (middle) and CTCF (right) across the mouse stem cell genome – indicative of dynamic binding trends seen for SETDB1 and Cohesin. **H**, Classification of annotated DiScs into functional genomic regions – simple repeats, ERVs, LINE, SINE and non-repeat regions, in order to distinguish DiSCs from other SETDB1-bound regions that are dominated by repeats. **I**, Heat map depicting the reproducibility of SETDB1 binding across SETDB1, H3K9me3 co-bound regions in public ChIP-Seq datasets as compared to our SETDB1 ChIP-Seq. **J**, Heat map depicting the reproducibility of SETDB1 binding across DiSC regions in public ChIP-Seq datasets as compared to our SETDB1 ChIP-Seq. **K**, Average enrichment profiles for SETDB1 across DiSCs as well as SETDB1, H3K9me3 co-bound sites, as compared between our ChIP-Seq and public SETDB1 ChIP-Seq (Bilodeau *et al.*, 2009). **L**, Spearman correlation plots comparing our ChIP-Seq to SETDB1 ChIP-Seq (Bilodeau *et al.*, 2009) (spearman=0.58), and to SETDB1 ChIP-Seq (Yuan *et al.*, 2009) (spearman=0.49). **M**, Overlap for SETDB1 peaks detected in two public datasets - (Bilodeau *et al.*, 2009) and (Yuan *et al.*, 2009), revealing low common peak numbers.

### Supplementary Figure 2

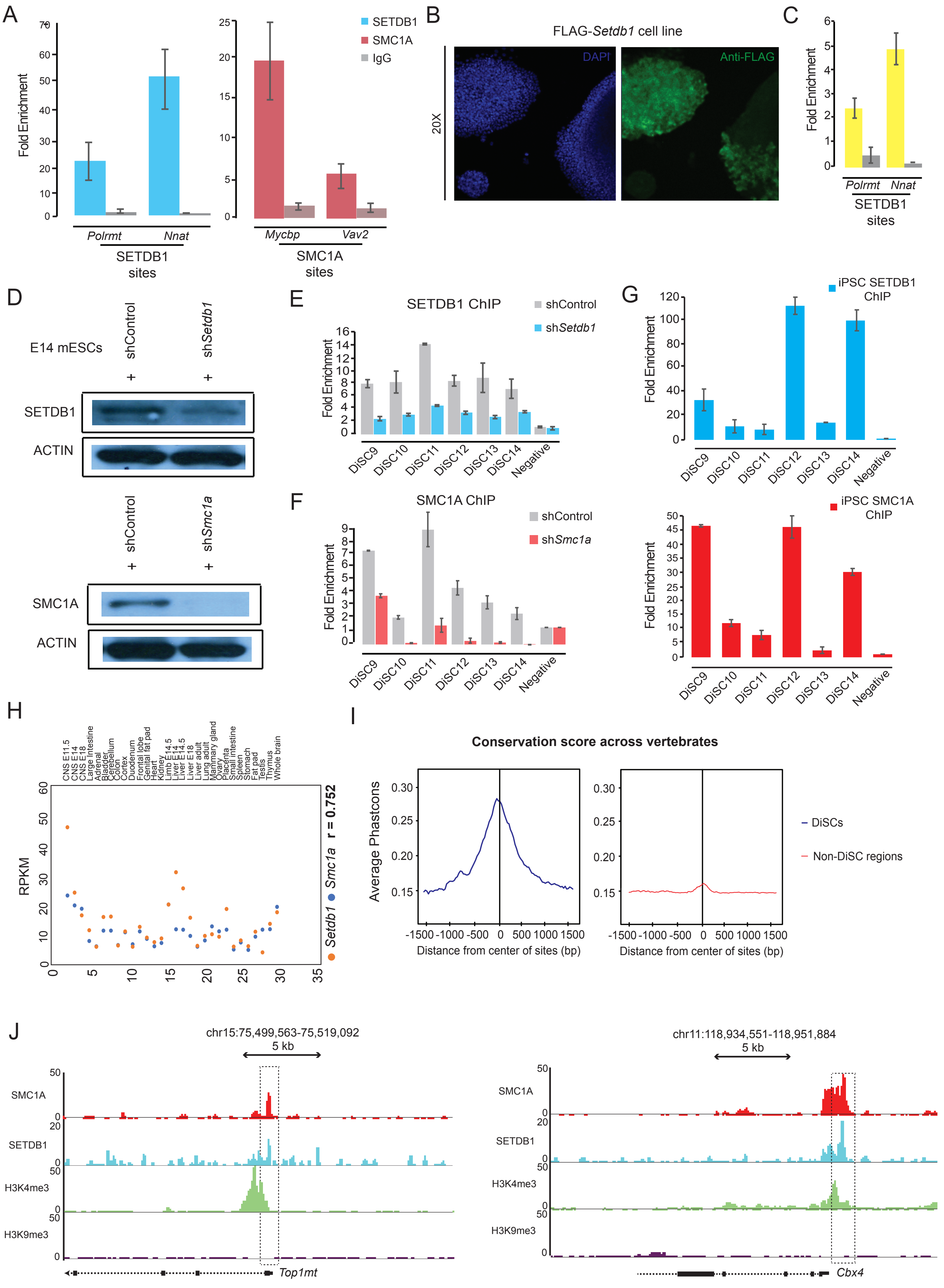

**Supplementary Fig.2 | SETDB1, Cohesin co-operation defines DiSCs and imparts them distinct and conserved features**. **A**, ChIP-qPCR validation of positive control sites for SETDB1 (left) and SMC1A (right) binding (n=3, Error bars represent standard deviation). **B**, Immunofluorescence-based validation of FLAG-*Setdb1* cell line displaying abundance of SETDB1 in the nucleus. **C**, ChIP-qPCR validation of positive control sites for SETDB1 using the FLAG-Setdb1 cell line. (n=2, Error bars represent standard deviation). **D**, Western blot depicting decreased protein levels for SETDB1 and SMC1A upon knock-down of these genes by shRNA constructs. **E**, KD of SETDB1 followed by SETDB1 ChIP-qPCR depicts the decline in SETDB1 binding at DiSCs. (n=3, Error bars represent standard deviation). **F**, KD of SMC1A followed by SMC1A ChIP-qPCR depicts the decline in SMC1A binding at DiSCs (n=3, Error bars represent standard deviation). **G**, ChIP-qPCR for SETDB1 and SMC1A representing their binding at DiSCs in iPSCs reprogrammed from mEF cells (n=3, Error bars represent standard deviation). **H**, Scatter plot representing correlation between the expression of *Setdb1* and *Smc1a* across different cell and tissue types derived from the ENCODE dataset. **I**, Average profiles illustrating PhastCon scores generated for DiSCs across vertebrate species. PhastCon scores represent conservation of specific sequences and their associated features. (PhastConDiSC ~ 0.28). An equal number of randomised genomic regions is chosen as background for this comparison. The PhastCon score generated was derived from a panel of 20 vertebrate species. **J**, UCSC browser screenshots representing the presence of the activating histone mark H3K4me3 proximal to the DiSCs.

### Supplementary Figure 3

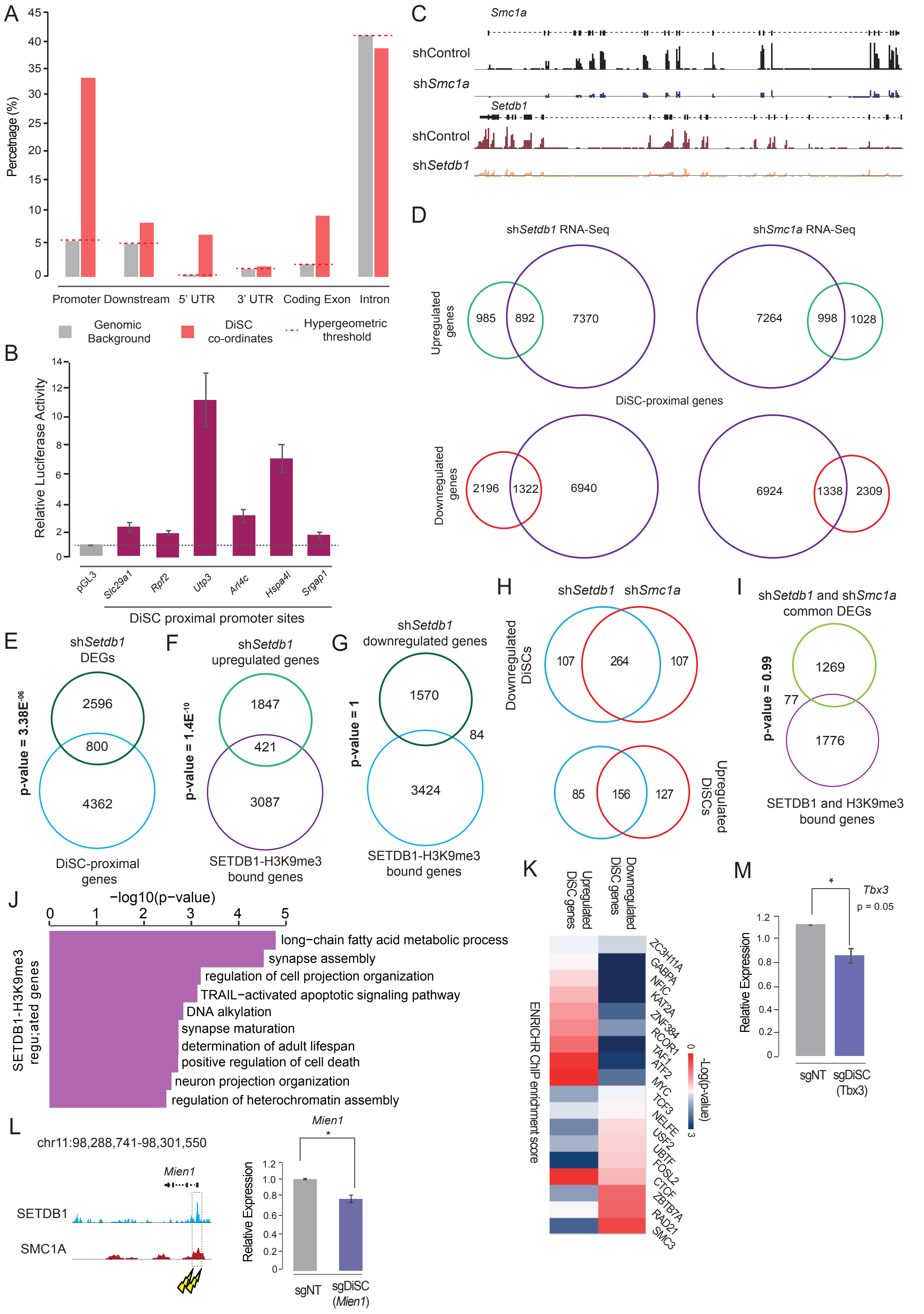

**Supplementary Fig.3 | DiSCs are critical genomic domains engaged in transcriptional regulation. A**, Hyper-geometric test indicating the distribution of DiSCs over different genomic regions recognising a prominent presence of these sites closer to promoters and the 5'UTRs of proximal genes. **B**, Luciferase assay for DiSC proximal gene promoters, indicating active transcriptional activity at these sites (n=2, Error bars represent standard deviation). **C**, UCSC browser-based representation for RNA-Seq libraries for sh*Setdb1* and sh*Smc1a* samples depicting the efficient KD of the targeted genes. **D**, Venn diagram depicting the DiSC-proximal genes that are upregulated and downregulated in sh*Setdb1* (left) *and* sh*Smc1a* (right) RNA-Seq samples. **E**, A stringent overlap between sh*Setdb1* DEGs and DiSC-proximal dysregulated genes revealed 800 such genes as common between the two sets ($p = 3.38E^{-06}$). **F**, Overlap between SETDB1, H3K9me3 co-bound genes and upregulated genes in sh*Setdb1* libraries ($p = 1.4E^{-10}$). **G**, Overlap between SETDB1, H3K9me3 co-bound genes and downregulated genes in sh*Setdb1* libraries (p = 1). **H**, Venn diagrams illustrating the genes that are commonly downregulated (top) or upregulated (bottom) by KD of SETDB1 and SMC1A, proximal to the DiSCs. **I**, Overlap between common DEGs in sh*Setdb1* and sh*Smc1a* treated cells and SETDB1, H3K9me3 co-bound genes (p = 0.99). **J**, Gene ontology (GO) analysis for SETDB1, H3K9me3 co-bound genes. **K**, Heat map indicating ENRICHR profiles for factors enriched at upregulated and downregulated DiSC genes, that were commonly regulated by SETDB1 and SMC1A. **L**, Quantitative PCR based validation (right) of *Mien1* downregulation after CRISPR-Cas9 knock-out of the SETDB1-SMC1A binding locus for the proximal DiSC (left) (n=3, Error bars represent standard deviation) [*p-value < 0.05, **p-value < 0.01]. **M**, Decrease in the expression of *Tbx3*, upon successful KO of the SETDB1-SMC1A binding locus of the proximal DiSC (n=3, Error bars represent standard deviation). [*p-value < 0.05, **p-value < 0.01].

### Supplementary Figure 4

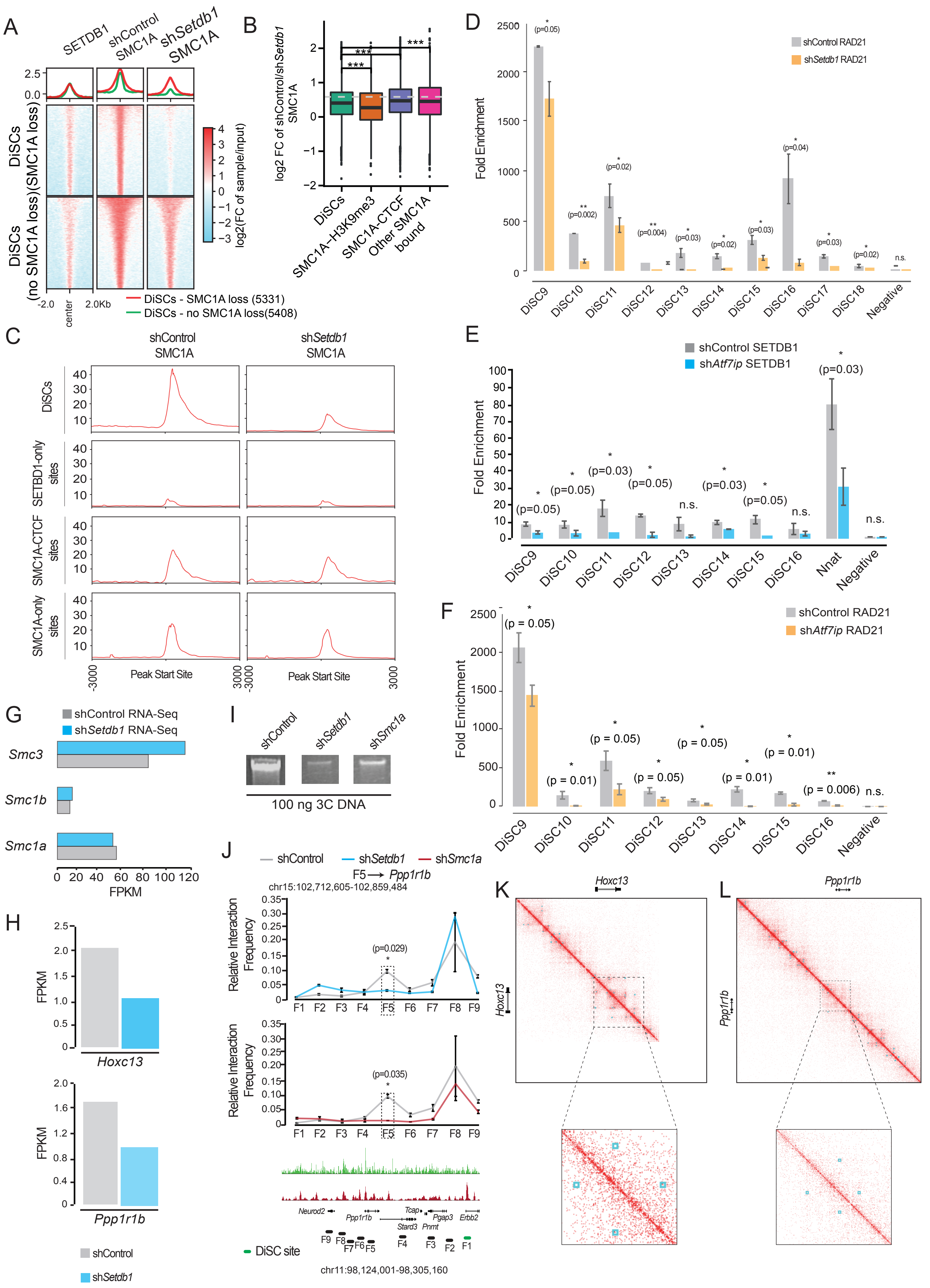

**Supplementary Fig.4 | SETDB1 functions as a core regulator of Cohesin-DNA associations.** **A**, Heat maps sub-categorizing DiSCs based on the decrease in SMC1A binding upon SETDB1 KD. **B**, Box-plot indicating the changes in SMC1A levels upon SETDB1 KD, across DiSCs, SETDB1-H3K9me3 bound sites, SMC1A-CTCF bound sites and other SMC1A-bound sites. **C**, Average enrichment profiles for SMC1A enrichment in shControl and sh*Setdb1* treated cells, representing the level of decrease of the same across DiSCs, SMC1A-CTCF bound regions, SETDB1-only and SMC1A-only bound sites. **D**, ChIP-qPCR data depicting decrease of RAD21 binding at DiSC sites upon KD of SETDB1. Values are mean + s.e.m for independent replicate experiments. Two-tailed t-test was used for statistical analysis. Error bars represent standard deviation (n = 3). [*p-value < 0.05, **p-value < 0.01]. **E**, ChIP-qPCR data depicting decrease of SETDB1 binding at DiSC sites upon KD of ATF7IP. Values are mean + s.e.m for independent replicate experiments. Two-tailed t-test was used for statistical analysis. Error bars represent standard deviation (n = 3). [*p-value < 0.05, **p-value < 0.01]. **F**, ChIP-qPCR data depicting decrease of RAD21 binding at DiSC sites upon KD of ATF7IP. Values are mean + s.e.m for independent replicate experiments. Two-tailed t-test was used for statistical analysis. Error bars represent standard deviation (n = 3). [*p-value < 0.05, **p-value < 0.01]. **G**, Unperturbed expression profiles for Cohesin sub-unit genes (*Smc1a*, *Smc1b*, *Smc3*) as seen in sh*Setdb1* treated cells. **H**, Downregulation of *Hoxc13* and *Ppp1r1b* genes proximal to DiSCs (represented by sh*Setdb1* RNA-Seq data). **I**, Overall intensity profiles of 3C libraries prepared from shControl, sh*Setdb1* and sh*Smc1a* cell, delineating a decreased intensity of interactions in the sh*Setdb1*, sh*Smc1a* libraries – as seem upon running 100 ng of each library on an agarose gel. **J**, 3C-qPCR interaction profiles around *Ppp1r1b* genomic locus which is located proximal to a DiSC, upon KD of SETDB1 (top) and SMC1A (bottom). (n=3, Error bars represent standard deviation). Two-tailed t-test was used for statistical analysis. [*p-value < 0.05, **p-value < 0.01]. **K**, shControl Hi-C data visualised using Juicebox at the *Hoxc13* locus that was used for 3C-qPCR. **L**, shControl Hi-C data visualised using Juicebox at the *Ppp1r1b* locus that was used for 3C-qPCR.

Supplementary Figure 5

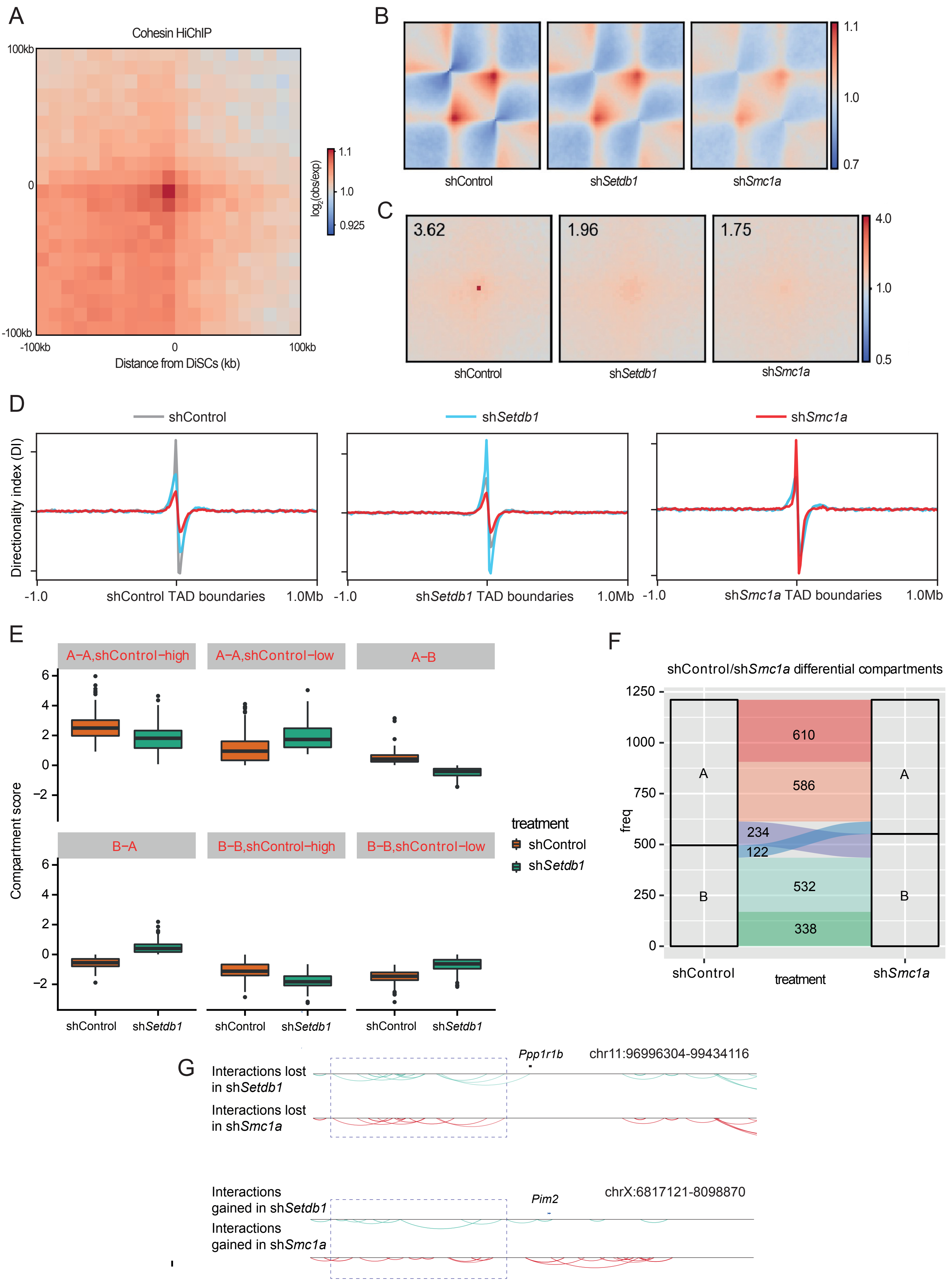

**Supplementary Fig.5 | SETDB1 is a key topological regulator aside from being an epigenetic modifier. A**, Overall enrichment plot for all DiSCs across Cohesin HiChIP data, at a 10 kb resolution. Normalization scale refers to logarithm (base-2) of observed versus expected enrichment. **B**, Enrichment plots displaying the genome-wide impact of sh*Setdb1* and sh*Smc1a* treatment on TADs, as compared to the shControl dataset (10 kb resolution). **C**, Genome-wide DNA loop enrichment in shControl, sh*Setdb1* and sh*Smc1a* Hi-C datasets, at a 5 kb resolution. **D**, Directionality Index (DI) profiles for shControl, sh*Setdb1* and sh*Smc1a* Hi-C data, indicating unique upstream/downstream preferential interactions for TAD/loop peripheries. **E**, Box-plot representation of all potential compartment transitions and compartment switches (A-B, B-A) between shControl and sh*Setdb1* treated cells. **F**, Transition plot indicating number of compartment switches arising between shControl and sh*Smc1a* treated cells. **G**, WashU browser-based visualization of interactions lost at the *Ppp1r1b* locus and its vicinity (top) and interactions gained at the *Pim2* locus and its vicinity (bottom) in sh*Setdb1* and sh*Smc1a* treated cells.
